# Urinary glucocorticoids in harbour seal (*Phoca vitulina*) pups during rehabilitation

**DOI:** 10.1101/549386

**Authors:** Susan C. Wilson, Stella Villanueva, Kayleigh A. Jones, William Hayes, Lilia Dmitrieva, Wesley Smyth

**Author notes:** Corresponding author: Susan C. Wilson, present address: 13 Wilmot Way, Banstead, Surrey, SM7 2PZ.

## Abstract

The glucocorticoid (GC) hormone cortisol is often measured in animals to indicate their welfare and stress levels. However, the levels of other naturally occurring GCs are usually overlooked. We aimed to investigate whether aspects of the care and conditions of harbour seal (*Phoca vitulina*) pups in rehabilitation centres are reflected in urinary concentrations of four endogenous GCs. Urine samples were collected non-invasively from pups taken in as “orphans” at five different rehabilitation centres: three on the Irish Sea and two in the southern North Sea. Concentrations of urinary cortisol, cortisone, prednisolone and prednisone were analysed by mass spectrometry. Urinary concentrations of endogenous prednisolone and prednisone occurred in similar magnitude to cortisol, for the first time in any mammal species. The levels of all GC concentrations decreased as pups gained mass, but the most significant effect was for prednisone. Pups with mass less than 11kg, i.e. healthy average birth mass, had significantly higher levels of prednisone (but not of the other GCs) than pups of 11kg or more. Cortisol, cortisone and prednisolone concentrations were slightly higher for pups without access to water than those with water; however, we found no significant effect of social group on GC levels. Based on these findings, we tentatively suggest that the GCs may be elevated in harbour seal pups during rehabilitation in response to some physiological factors deviating from the norm of free-living pups. Our findings highlight the importance of measuring other GCs, in addition to cortisol, for understanding stressors affecting the welfare of seal pup in rehabilitation.

## 1. INTRODUCTION

New-born harbour seal pups found stranded on the shoreline without the mother and believed to be permanently separated from her (termed ‘orphan’ pups; see Wilson and Jones, 2021) are typically taken to ‘seal sanctuary’ centres for mother-substitute care (usually termed ‘rehabilitation’) until they are considered well-enough grown to be released back into the sea, usually after a period varying between about seven weeks to 4 months.

The method of handling and feeding these orphan pups, the nutritional content of their food and both their social and physical environment varies widely between different rehabilitation centres. However, rehab centres are only able to provide the pups with an artificial environment which differs in many key aspects from the natural environment of mother-dependent, free-ranging pups. It is therefore likely that the rehab environment exposes the pups to many stressors that differ from the stressors in the pups’ natural environment. Gulland et al. (1999) found changes in adrenal function, manifest as decreased cortisol production following injection of adrenocorticotrophic hormone (ACTH), in pups surviving eight weeks of rehabilitation. These authors suggested this might be due either to adaptation to captivity or to adrenal exhaustion.

Mother-dependent, free-ranging pups spend around 60% of their time in the water from birth (Bowen et al., 1999; Skinner, 2006). Average birth weight is about 11 kg and pups nurse from their mother usually for 3–4 weeks, or up to 6 weeks in some populations, with pups typically achieving an average net weight gain of 0.4 to 0.6 kg/day (Skinner, 2006; Muelbert and Bowen, 1993; Cottrell et al., 2002). The fat content of the milk increases from about 41% at birth to 50% by day 7 until weaning (Lang et al., 2005). During the nursing period, free-living pups are constantly close to their mother (Venables and Venables, 1955; Wilson, 1974; Wilson and Jones, 2018) until about 10 days of age, when mothers sometimes leave their pups onshore in the haul-out group for a few hours while they make foraging trips offshore (Wilson, 1978; Boness et al., 1994). Reproducing – for orphan pups in rehab - this concentrated nutrition and key features of the amphibious and social environment of free-living pups is obviously challenging.

Harbour seal pups in rehab will usually not suckle from a bottle. Some centres feed pups by force-feeding with whole fish or tube-feeding fish ‘soup’, with little or no growth or weight gain by pups until they are past natural weaning age and feeding freely on whole fish (Wilson et al., 1999; MacRae et al., 2011). Other centres enhance pup weight gain by tube-feeding pups enriched formulae using dedicated high fat, multi-milk-based formulations; however, average weight gain nevertheless rarely achieves 0.3 kg/day (Robinson, 1995; Wilson, 1999; MacRae et al. 2011).

Rehab facilities in the British Isles and N. America are usually designed to maintain orphan harbour seal pups in isolation and in dry pens for some weeks after admission (Larmour 1989; Robinson, 1995; MacRae et al. 2011), a principal reason for this being to prevent potential transmission of infections (Osinga and ‘t Hart. 2010). Less often, centres house these pups with access to a pool, either in pairs (Wilson, 1999) or in small groups (Müller et al., 2003), where they can experience species-typical sensory stimulation and muscle movement during swimming and social interaction.

Trumble et al. (2013) considered cortisol to be the primary circulating corticosteroid in seals. These authors investigated serum cortisol levels in free-ranging and rehab harbour seal pups. The cortisol levels increased in pups during their first weeks in rehab while tube-fed fed on milk formula, but were lower in pups that were fish-fed after eight weeks of age. Body mass was not a significant predictor of cortisol levels, although the lower cortisol levels in the older, fish-fed pups were similar to free-ranging pups. However, the social and physical environments of the rehab pups during milk and fish feeding were not specified in this analysis (Trumble et al. 2013).

When a dependent, free-living harbour seal pup is separated from its mother, it presumably experiences acute stress. Serum cortisol levels were found to be elevated in pups when they were captured alone, but the levels were lower if the mother was with the pup during capture (Di Poi et al., 2015), suggesting that acute stress was avoided if the pup was not separated from its mother.

When a free-living harbour seal pup strands due to having been permanently separated from its mother, it may be taken to a rehabilitation centre, where it is fed and cared for until it is ready for release back to the sea. By the time an “orphan” pup enters rehab, it is likely past the initial “acute” stress stage of searching for its mother and may already be in the next “depressive” phase of maternal separation, when it may be expected to display more passive behaviours (Spencer-Booth and Hinde, 1971) accompanied by reduced circulating cortisol (Yusko et al, 2012).

The conditions for the first few weeks in rehab vary according to centre, from social isolation in a dry pen (Robinson, 1995; Osinga and ‘t Hart, 2010) to a pair or group with water access (Wilson 1999), and from feeding fish or low-calorie fish “soup” to higher calorie milk formulations (Macrae et al., 2011, Wilson, 1999). However, it is not known how these varied conditions in different rehabilitation centres may affect glucocorticoid hormone levels. Elevated cortisol levels have often been thought to reflect poor welfare of captive animals, although this has been questioned and the cortisol-welfare relationship is complex, depending on the context (e.g. Cook, 2000; Linklater et al. 2010; Novak et al., 2013; Otovic and Hutchinson, 2014).

It seems most likely that orphan pups kept socially with free water access would maximise their psychological welfare, because these conditions allow pups to express many of the natural social and physical behaviours of free-living pups with their mothers (Wilson, 1999; Wilson and Jones, 2018; S. Wilson and R. Alger, unpublished data). Our initial hypothesis for this study was that orphan pups held in isolation in a dry pen would have elevated cortisol levels relative to orphans maintained socially with water access. The initial phase of the present study (2012–2013) therefore explored cortisol levels in two rehabilitation centres with those contrasting conditions. However, when the opportunity arose (in 2014), we expanded the study to include pups in further rehabilitation centres and to consider all four potentially endogenous glucocorticoid hormones, i.e. cortisone, prednisolone and prednisone, in addition to cortisol. Our correspondingly revised hypothesis was that different components of the pups’ rehabilitation conditions (i.e. the social or physical environment and nutritional state) would be reflected in the levels of one or more of these four GC hormones.

Our study differed from previous studies in harbour seals in that we chose to sample urine rather than blood, since urine sample collection would be non-invasive, and would reflect GC hormone concentrations over time since the previous voiding (Cook, 2012, Novak et al., 2013). The present study is exploratory and purely descriptive, since aspects of the care and conditions of pups at each centre could not, for ethical reasons, be manipulated for research purposes.

## 2. ANIMALS AND METHODS

### 2.1 Pup background

This study involved harbour seal pups that had been admitted to five participating rehabilitation centres in Europe. All the pups had been found stranded on the shoreline during the pup birthing season. On admittance to the centres, the pups were mostly at or below average birth weight of approximately 11kg for harbour seals. They were therefore all believed to be ‘orphans’, having been permanently separated from their mothers in the neonatal or early post-natal period. None of the pups were of sufficient size to have been healthy pups, mistakenly taken into captivity while their mothers had left them onshore temporarily while foraging offshore (Boness et al., 1994; Wilson and Jones, 2021). No pups in the study received treatment with synthetic steroids.

### 2.2 Pup rehabilitation centres

Five seal pup rehabilitation centres participated in this study. Two were on the southern North Sea coast: centre I in Lincolnshire, UK and centre IV in northern Germany. The other three centres (II, III and V) were located on the Irish Sea coast of Ireland. There were therefore differences among the populations in which pups were born into, in addition to differences in pup care procedures and rehab captive environments in the centres (Table 1) during the sampling period. In centres I and III, the pups were kept alone in dry pens, in centres II and IV the pups were usually in pairs or groups with free access to water, and in centre V the pups were either alone or paired and in dry pens (Fig. 1; Table 1). The water in centres II and IV was sufficient for the pups to submerge while socialising or sleeping (Fig. 2).

**Fig. 1.**
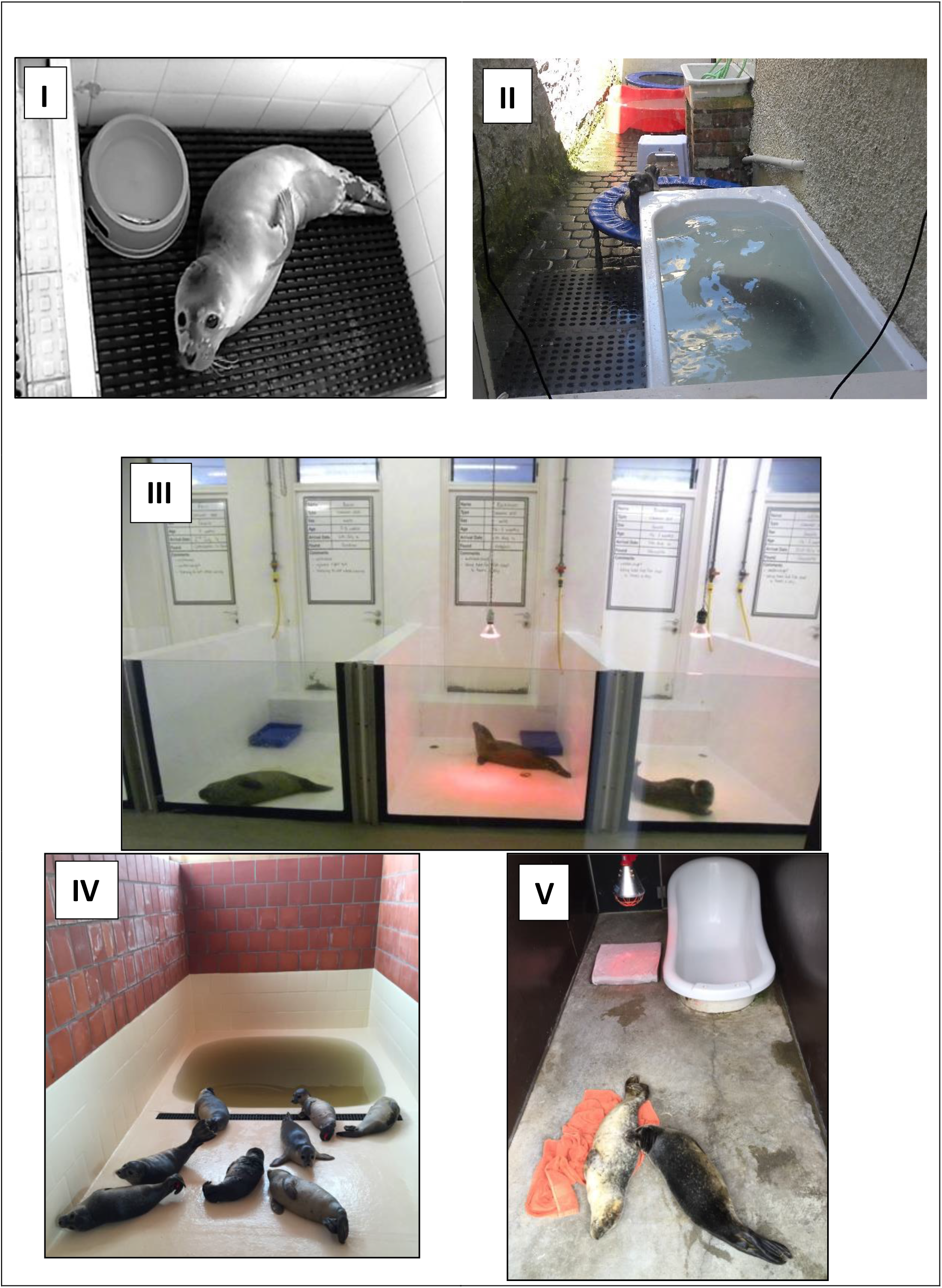
Pups at each of the five centres. **I** – indoor pens ~110×80 or 110×160 cm, heating lamp; **II** – outdoor open enclosure ~130×600 cm including two filled pools or pool and filled bath and two trampolines, no heating lamp; **III** – indoor pens ~100×200 cm, heating lamp; **IV** – outdoor covered enclosures, varying dimensions, with pool and heating lamp; **V** – outdoor covered enclosure ~ 120×320 cm, bath filled only for fish-feeding, heating lamp.

**Fig. 2.**
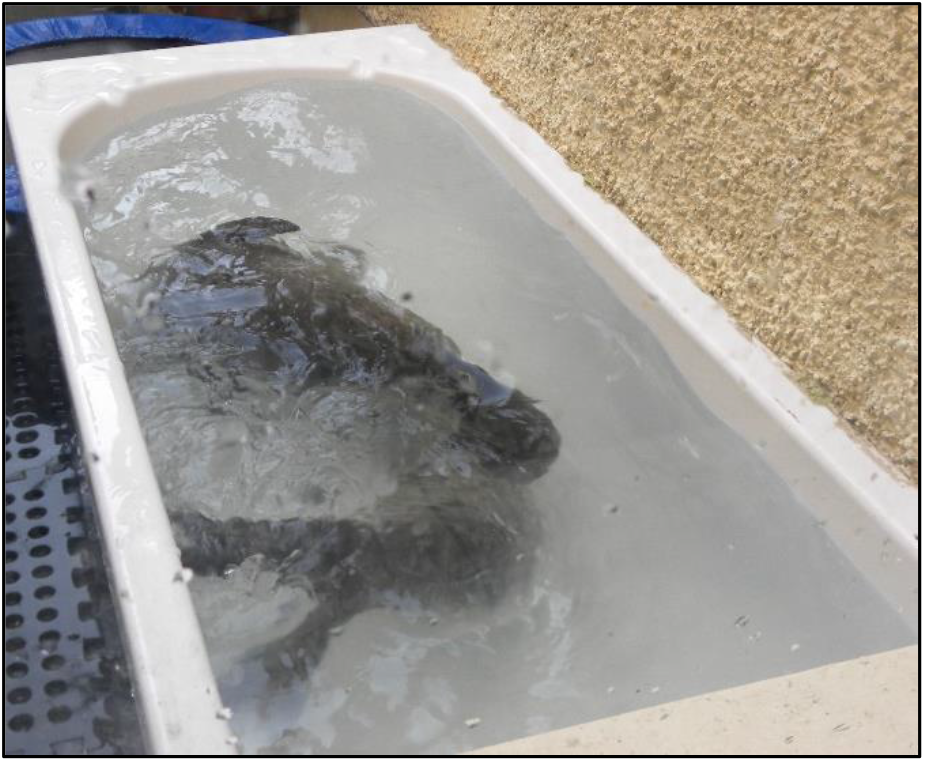
Paired pups (rehab days 16 and 18) in centre II playing together while submerged in bath.

**Table 1.** Number of pups at each of five rehab centres that were urine-sampled in the study, with summary of rehab environment and average growth rate (kg/day) at each centre.

| **Centre** | **I** | **II** | **III** | **IV** | **V** |
| --- | --- | --- | --- | --- | --- |
| No. pups sampled | 3 | 6 | 7 | 13 | 6 |
| Sex M/F | 3M | 2M/4F | 5M/2F | 6M/7F | 4M/2F |
| Pups' food | whole herring | *Pet Ag* multimilk + fish + mixed fish oil | herring soup + salmon oil | herring soup and/or whole herring | herring soup + salmon oil and/or whole herring |
| Est. Pup mass (kg) on sample day | 9.7 | 15.6 | 9.4 | 11.6 | 11.4 |
| (range) | (9.5–10.3) | (9.3–21.5) | (7.9–11.8) | (7.1–14.9) | (9.2–14.9) |
| (SD) | 0.31 | 4.14 | 1.48 | 1.74 | 1.82 |
| Growth rate (kg/day) | 0.06 | 0.29 | 0.21 | 0.15 | 0.07 |
| (range) | (0.03–0.08) | (0.27–0.30) | (0.1–0.26) | (0.01–0.26) | (-0.1–0.19) |
| (SD) | 0.04 | 0.01 | 0.06 | 0.07 | 0.10 |
| Social groups at each centre | 1 | 2 | 1 | 1, 2, 3 | 1, 2 |
| Free water access | No | Yes | No | Yes | No |
| Rehab days (range) of sampling period | 13 (4–21) | 25 (8–44) | 6 (2–12) | 6 (2–17) | 19 (7–40) |

Pups were weighed routinely by centre personnel on alternate days at centre II and approximately once per week at the other centres. Average growth rates (kg/day) during the sampling period were calculated from the weighings at the beginning and end of the sampling period. Estimated pup mass on the sampling day was calculated as the mid-point between two adjacent weighings in centre II, and in the other centres from the nearest pup weighing and average growth rate during the sampling period.

### 2.3 Urine sampling

This study used urine rather than blood for the sampling to be completely non-invasive. The urine samples were collected opportunistically if the pups urinated while being handled during routine feeding or weighing. The authors (SW and SV) collected samples at centres II, IV and V, while centre personnel collected samples at centres I and III. Collection at centres II–V was usually with a syringe from the ground beside the pup or directly from the stream of urine, although one sample from centre II was collected on a ‘puppy pad’, with plastic-backed paper tissue layers. Most samples at centre I were collected from a tray under the slatted floor of the pen during cleaning (samples collected in the morning were therefore from urine voided at any time since the previous evening). Samples were transferred to standard plastic sample pots with a label detailing the source pup, date and collection time. Samples were then immediately transferred to a freezer (at least −18°C) and stored until analysis. Only samples with at least 10 ml were used in the analyses.

It has been suggested, in bovines, that neo-formation of the GC prednisolone may occur due to microbial action after urine collection if the samples are kept at a high temperature (e.g. 37°) or if there is contamination (e.g. from faeces) (Arioli et al. 2012). We were therefore careful to ensure that the samples used for analysis were free from visible faecal contamination, and were not subjected to high temperatures.

Since a study of plasma cortisol levels in captive harbour seals found a diurnal pattern, peaking at 01:00 and waning around 13:00 (Gardiner & Hall 2012), the time of day of each sample collection was recorded and samples from pup rehab day 2 onwards (n=112) were assigned to time categories: 0700–12:59; 13:00–18:59; 19:00–00:59. No samples were collected during the overnight period.

Seven samples were collected from a pair of pups (in Centres II and IV), but the donor pup was not identified (Supporting file S1). These samples were included in analyses of the overall range of GC concentrations but were excluded from subsequent analyses.

### 2.4 Analysis of urinary glucocorticoid concentrations

If the concentrations of several steroids are required, either separate kits for each steroid must be used (requiring splitting of the sample), or the steroids need to be separated by chromatography. Ultra-Performance Liquid Chromatography (UPLC) coupled with tandem mass spectrometry (UPLC-MS/MS) can quantify either one or multiple steroids at wide-ranging concentrations, while avoiding the issue of antibody cross-reactivity (Koren et al. 2012). The UPLC-MS/MS methodology used in the present study to analyse urine samples from seal pups has been described in detail by McWhinney et al., 2010, De Clercq et al., 2013 and reviewed by Hawley and Keevil (2016).

Eleven samples from Centre 1 (2012) and eight from centre II (2013) were analysed for CL at the Institute of Global Food Research, Queens University, Belfast (Lab 1). Ninety-eight further samples collected from centres II, III, IV and V in 2014–16 were analysed at the Chemical and Immunodiagnostic Sciences Branch, Veterinary Sciences Division, Agri-Food & Biosciences Institute (AFBI), Belfast (Lab 2) for CL, cortisone (CN), prednisolone (PL) and prednisone (PN).

The CL analyses of the 2012-2013 samples were carried out alongside a separate study of CL and CN concentrations in cattle in Lab 1 (Keenan, 2015), while the analyses of CL, CN, PL and PN in the 2014–2016 samples were carried out according to the procedures for analysis of bovine liver as part of the routine statutory Veterinary Drug Residue testing programme in N. Ireland (Lab 2).

The UPLC-MS/MS protocols followed by each laboratory are detailed in supplementary file S1.

The UPLC-MS/MS protocols followed by each laboratory are detailed in supplementary file S1. In addition, a comparison of the results from 11 samples for cortisol concentrations in the two pups from centre 1 (in 2012) analysed by both UPLC-MS/MS and an ELISA immunoassay kit are presented separately in Supplementary file S2.

UPLC-MS/MS method validation for the 2012–2013 samples was previously undertaken for analysis accuracy/recovery, intra- and inter-day repeatability and linearity for bovine urine in Lab 1 (Keenan, 2015). Method validation in Lab 2 was previously carried out for bovine, porcine and ovine liver in accordance with EU guidelines (2002/657/EC).

#### 2.4.1 Urinary creatinine concentrations

To control for the water content of the sample, urinary GC concentrations are expressed as a ratio to the creatinine (Cr) concentration (ng GC mg/Cr) in each sample. Creatinine concentrations at both laboratories were analysed using a standard colorimetric method modified from the Jaffe reaction, in which Cr in an alkaline solution reacts with picric acid to form a coloured complex (Taussky and Kurzmann, 1954).

### 2.5 Statistical analyses

#### 2.5.1 Non-parametric tests

Non-parametric statistics tests were carried out in XLSTAT (Addinsoft, 2019). The concentrations of all four GCs for all samples were not normally distributed. The correlations between concentrations of the four GCs (CL, CN, PL and PN) in all samples were therefore assessed using non-parametric Spearman correlation. Some pups were sampled more than once during the sampling period, a median value of GC concentrations from each of these pups was obtained.

Since pups’ GC concentrations when they first arrive at the centre are likely to reflect their stranding conditions, handling and transport to the centre, single or median concentrations in pups were compared between days 0-1 and day 2 onwards using a Mann Whitney U test. Only samples from day 2 onwards were used in further analyses.

To compare CL and PN concentrations among different time periods of the day at each of the centres, Kruskal-Wallis 3-sample tests and Mann-Whitney 2-sample tests were used. Kruskal-Wallis tests were also used to determine whether GC concentrations significantly differed among the five centres, as well as to determine whether pup growth rate differed among the five centres. We also considered the possibility that pup mass at entry to the rehab centre may have affected its subsequent growth rate at the centre and tested this by Spearman correlation.

To investigate whether low body mass is associated with relatively high GCs, we compared GC concentrations in pups less than 11 kg (approx. average birth mass) with pups of mass ≥ 11 kg on the dates of urine sampling using a Mann Whitney U test.

#### 2.5.2 Generalised linear models

To determine which factors best explained GC concentrations in harbour seal pup urine, we ran linear mixed effects models for each GC in R (version 3.6.0) (R Core Team 2020), using the R package ‘nlme’ (Pinheiro et al. 2019). We used all the data from rehab day 2 onwards and we transformed the GC concentrations to achieve normal distributions of data: CL and CN concentrations were log-transformed, and PL and PN concentrations were square root transformed (denoted by √). Predictor variables in candidate models included sex, estimated mass on sample day, access to water, housing with or without other pups, as well as the interactions among these factors. We included individual pup as a random effect to account for variability among individuals. We selected the best-fit model for each GC according to the lowest AIC or the simplest model if models differed by less than 2 AIC. We did not include rehab day in this model as it was highly correlated with pup mass on sample day (correlation = 0.81, variance inflation factor = 5.8). However, we also separately ran the models with rehab day instead of pup mass.

## 3. RESULTS

### 3.1 Ranges, median concentrations and correlation matrix of GCs

The creatinine (Cr) concentrations in all the samples analysed by MS/MS ranged from 115–15576 μM/L (mean 3219, median 2273 μM/L). The GC concentrations (ng/mg Cr) were highest for CL (range 0–8062, mean 119, SD 846, median 36.4 ng/mg), followed by CN (0–4853, mean 526, SD 617, median 390 ng/mg), then PL (0–236.6, mean 39.9, SD 43.6, median 30.4 ng/mg) and PN (0–142.5, mean 36.7, SD 26.7, median 31.7 ng/mg). The median values for CN were therefore about ten times higher than for CL, PN and PL (Fig. 3). However, CL values were highly variable, with 3 of the 117 samples (all taken between rehab days 0–2) having values >2,000 ng/mg Cr. There were seven samples with undetectable CL and PL concentrations (i.e. zero ng/mg). Six of these samples had low Cr concentrations (ranging from 115–683 μM/L).

**Fig 3.**
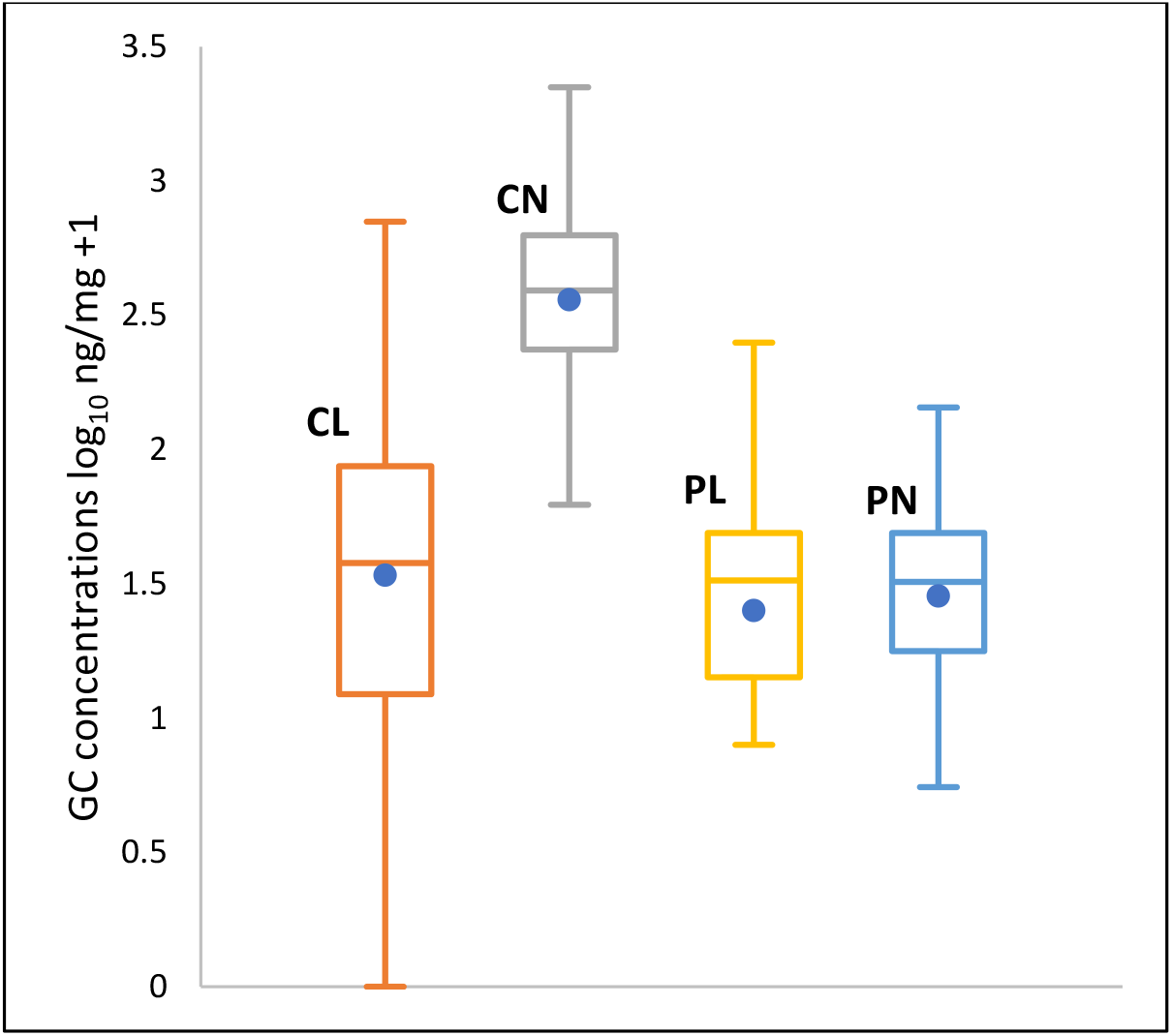
Box plots of urinary concentrations of four GCs from all samples analysed by UPLC MS/MS. (n=CL (Cortisol) –117; CN (Cortisone), PL (Prednisolone), PN (Prednisone) – 98). Concentrations in ng/mg creatinine on log10 scale. Horizontal lines indicate 1^st^ quartile, median and 3^rd^ quartile and spot markers indicate means.

The Spearman correlation matrix (Table 2) for all four GCs for the samples indicates a significant (P < 0.05) positive monotonal relationship between urinary concentrations all four GCs; however, the strongest correlation was between CL and PL, while the weakest was between CN and PL.

**Table 2.**
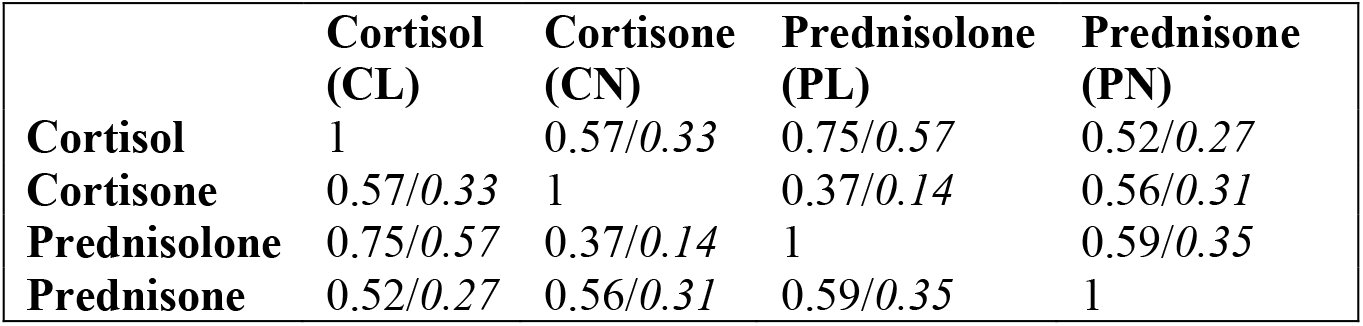
Spearman correlation matrix with correlation coefficients ρ between concentrations of four GCs (ng/mg creatinine) (n=98 samples; ρ *in italics*).

The ratios of median cortisol concentrations to the other three GCs from all samples from which all four GCs were analysed were CL:CN 0.09, CL:PL 0.18 and CL:PN 1.13, i.e. the ratios of median concentrations of CL to PL and PN were close to parity.

### 3.2 Comparison of GC concentrations of newly-admitted pups with those from rehab day 2 onwards

In days 0-1, eight pups were sampled for CL, and six of these pups were sampled for PN, CN and PL, and in day 2 onwards 34 pups were sampled for CL, PN, CN and PL. The concentrations of CL, PN and PL were significantly higher in days 0-1 than day 2 onwards (Mann-Whitney U tests; 1-tailed P <0.05 for each GC). However, CN concentrations did not significantly differ between days 0-1 and day 2 onwards (P = 0.328).

### 3.3 Effect of time of day of sampling on GC concentrations

The distribution of sampling from rehab day 2 onwards indicated that most of the samples were taken in the morning 6-hour period (0:700–12:59) and afternoon (13:00–18:59), with fewer in the evening (19:00–00:59). Concentrations of cortisol and prednisone, as representative GCs, were compared in each of the time periods in each centre. No significant differences between the time periods were found (Table 3).

**Table 3.** Median concentrations of urinary cortisol and prednisone (ng/mg creatinine) and significance of differences between each of three 6-hour time periods between 07:00 and 00:59. (samples from rehab day 2 onwards). P values from Kruskal-Wallis 3-sample test or Mann-Whitney 2-sample test; number of samples (*in brackets*).

| Time period | 07:00–12:59 | 13:00–18:59 | 19:00–00:59 |  |
| --- | --- | --- | --- | --- |
| CENTRE | Median concentration ng/mg creatinine |  |  | P value |
| Cortisol |  |  |  |  |
| I | 79.07 (6) | 62.83 (4) | zero samples | 0.931 |
| II | 22.27 (6) | 11.50 (20) | 15.78 (9) | 0.258 |
| III | 13.85 (3) | 0.95 (3) | 11.73 (3) | N/A |
| IV | 82.6 (11) | 298.38 (17) | 87.17 (5) | 0.994 |
| V | 27.67 (9) | 45.01 (6) | zero samples | 0.859 |
| Prednisone |  |  |  |  |
| II | 26.88 (5) | 11.55 (20) | 8.72 (8) | 0.091 |
| III | 54.13 (3) | 10.61 (3) | 37.12 (3) | N/A |
| IV | 41.52 (11) | 36.29 (17) | 34.85 (5) | 0.426 |
| V | 41.97 (9) | 49.55 (6) | zero samples | 0.680 |

### 3.4 Differences in pup GC concentrations between the five centres

The distribution of GC concentrations (median or single value for each pup) analysed by UPLC-MS/MS is shown in Fig. 4. Kruskal-Wallis tests indicated that concentrations of all four GCs differed significantly between centres (P < 0.0001 for each GC). Centre IV samples had overall the highest CL, CN and PL concentrations, whereas Centre II had the lowest CL, PN and PL concentrations, while Centre III had the lowest CL concentration.

**Fig. 4.**
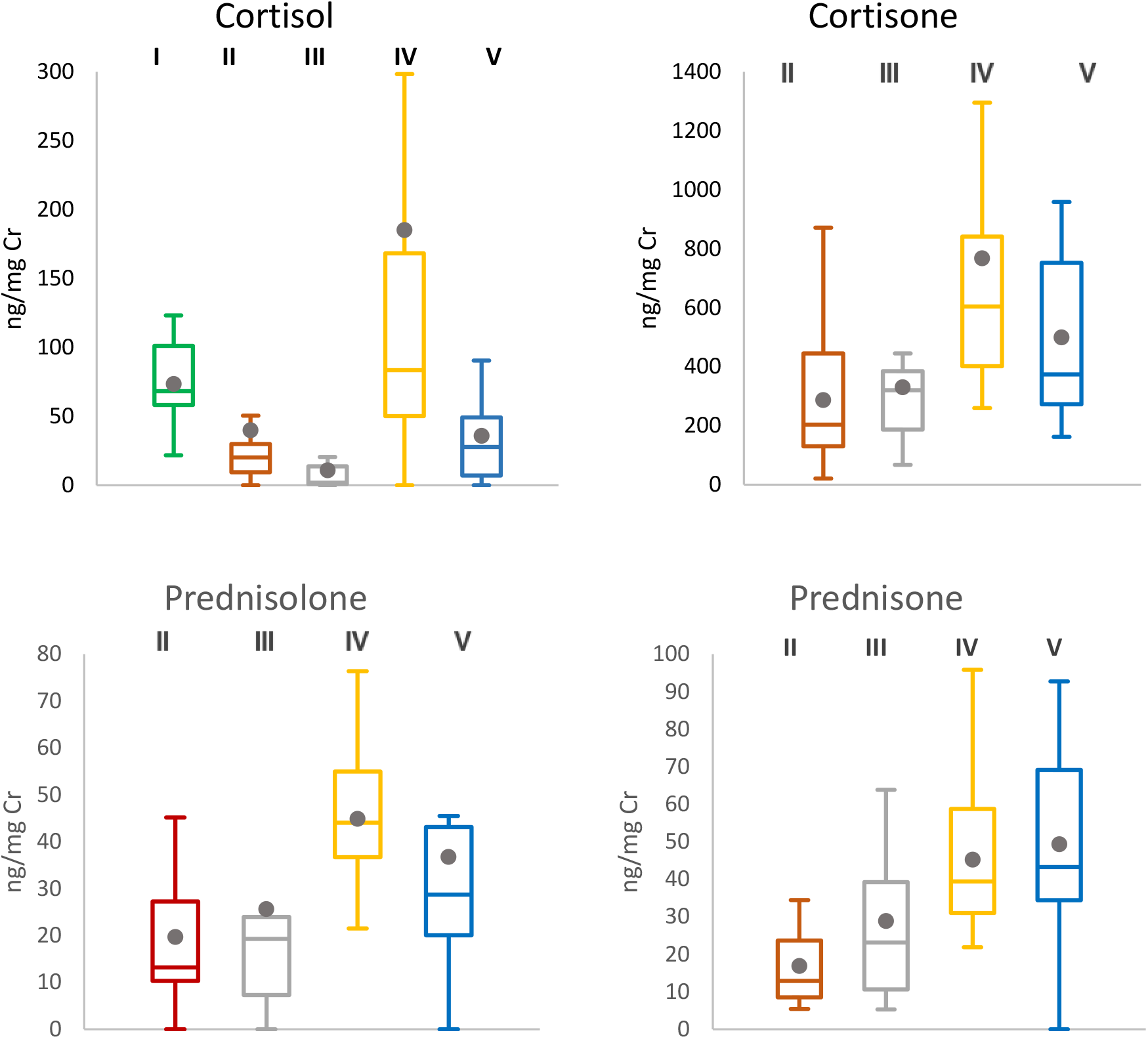
Boxplots of urinary concentrations of four GCs at each rehab centre I–V (day2 onwards), using individual and median values of GC concentrations for each pup as data points. Horizontal lines indicate 1^st^ quartile, median and 3^rd^ quartile and dot markers indicate means.

### 3.5 Pup growth rates, mass, and correlations with GC concentrations

There was a significant difference in pup growth rates between the five centres (Table 1; Kruskal-Wallis test, P <0.001). However, there was no apparent association between entry mass and individual pup growth rate during the sampling period (Spearman correlation coefficient ρ = 0.02). There was a significant negative monotonal relationship between individual growth rate and PL concentrations (Spearman coefficient ρ=0.372, P<0.025), but the negative relationship was not significant at the 5% level for the other GCs (Fig. 5).

**Fig. 5.**
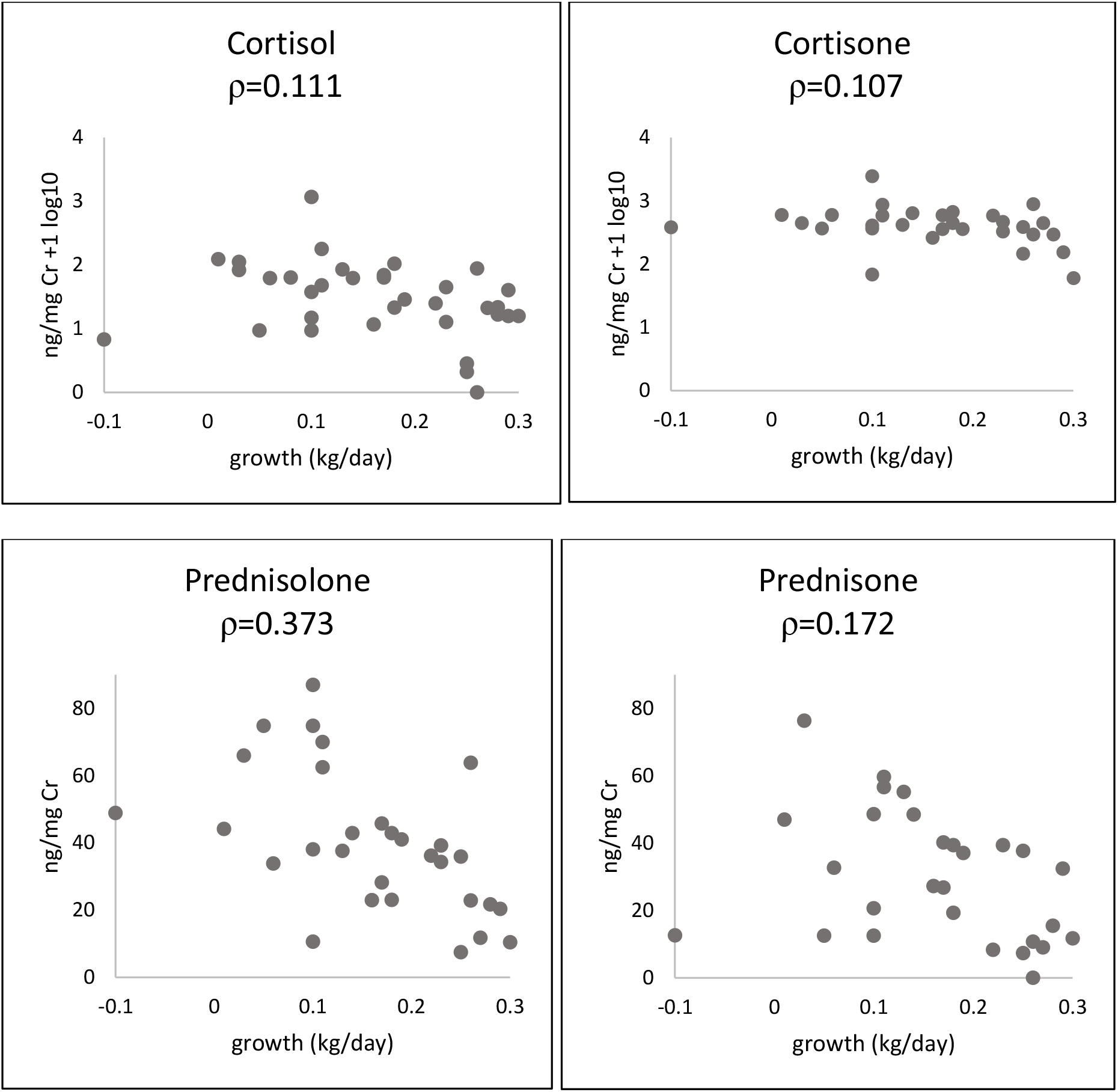
Scatter plots showing monotonal relationship, with Spearman correlation coefficient ρ, between median GC concentrations and average growth rate (during the sample period) for each pup sampled from rehab day 2 on. Plots for cortisol and cortisone are given as log10 of the concentrations +1, owing to some zero concentrations for cortisol and relatively high concentrations of cortisone.

A comparison of the range of concentrations of each GC for pups <11 kg (i.e. less than average healthy birth mass) and pups ≥ 11kg suggests that the range of concentrations was generally greater for the smaller pups (Fig. 6). The PN concentrations were significantly higher for pups <11 kg than for pups ≥ 11kg (P=0.009 Mann-Whitney U test) but there was no significant difference for the other GCs.

**Fig. 6.**
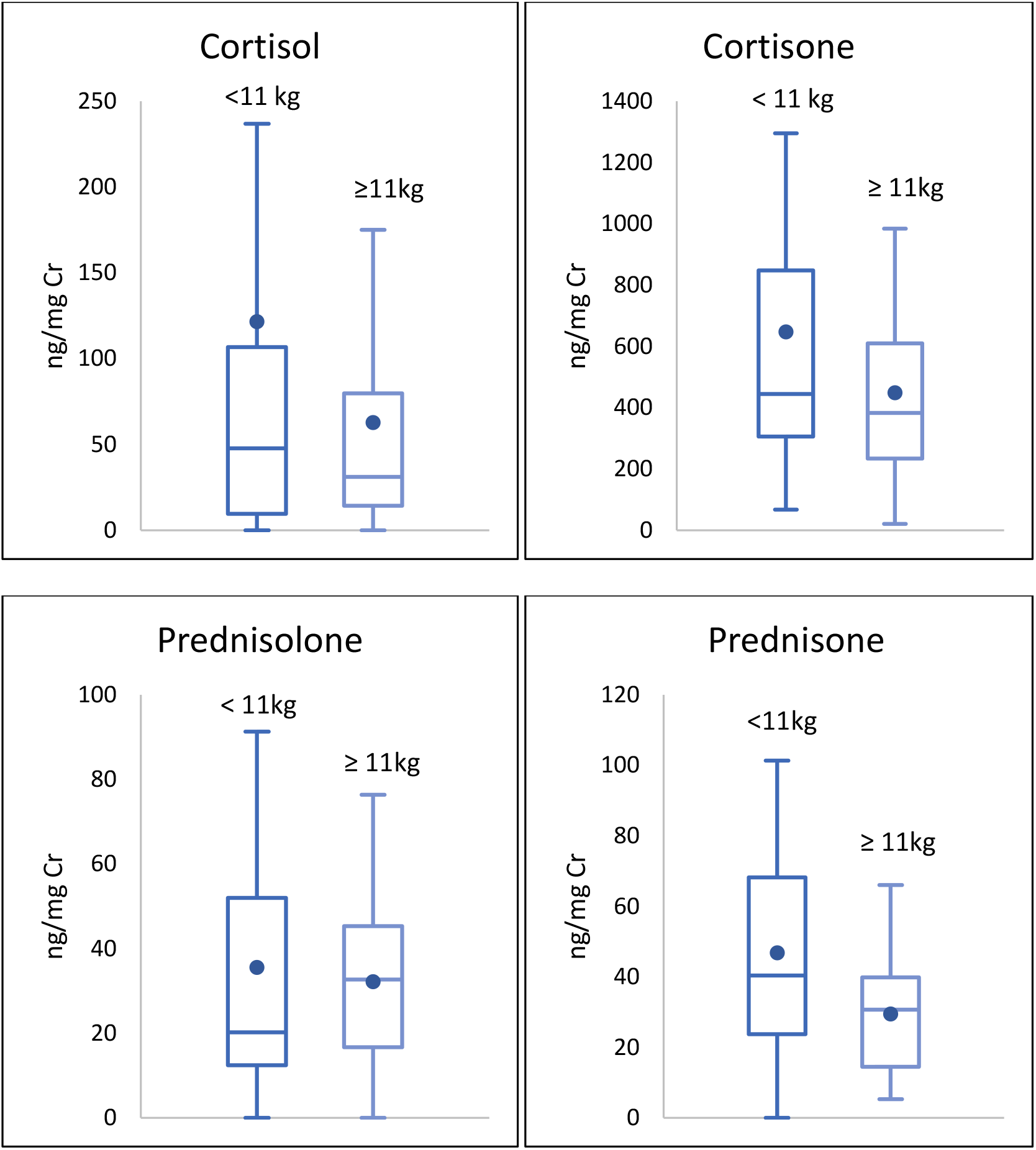
Urinary GC concentrations (ng/mg Cr) for pups (rehab day 2 onwards) weighing < 11 kg and ≥ 11 kg on date of urine sampling (outliers are not shown). Horizontal line indicates median, dot indicates mean. For pups < 11 kg, n= 37 samples from 17 individual pups for CL and 27 samples from 15 individual pups for CN, PL and PN. For pups ≥11 kg, n= 59 samples from 22 individual pups for CL and 51 samples from 20 pups for CN, PL and PN.

### 3.6 Predictor variables influencing urinary GC concentrations

Linear mixed effects models indicated that neither pup sex nor social group (alone or social) had a significant influence on concentrations of any of the urinary GCs. The predictor variables that best explained CL and CN concentrations were pup mass on sampling day, whether the pup had access to water, and the interaction between these factors (Table 4; Fig 7a, b). As pup mass increased, these urinary GC concentrations declined for pups with water access, but increased for pups without water access (Fig. 7a, b). The predictor variables that best explained PL concentration were pup mass on sampling day and whether the pup had access to water, with pups with water having slightly higher PL concentrations than those without (Fig. 7c). However, the second best-fit model (√prednisolone-2) was within 2 AIC of the first and included an interaction between these two factors (Fig. 7c-2).

**Fig. 7.**
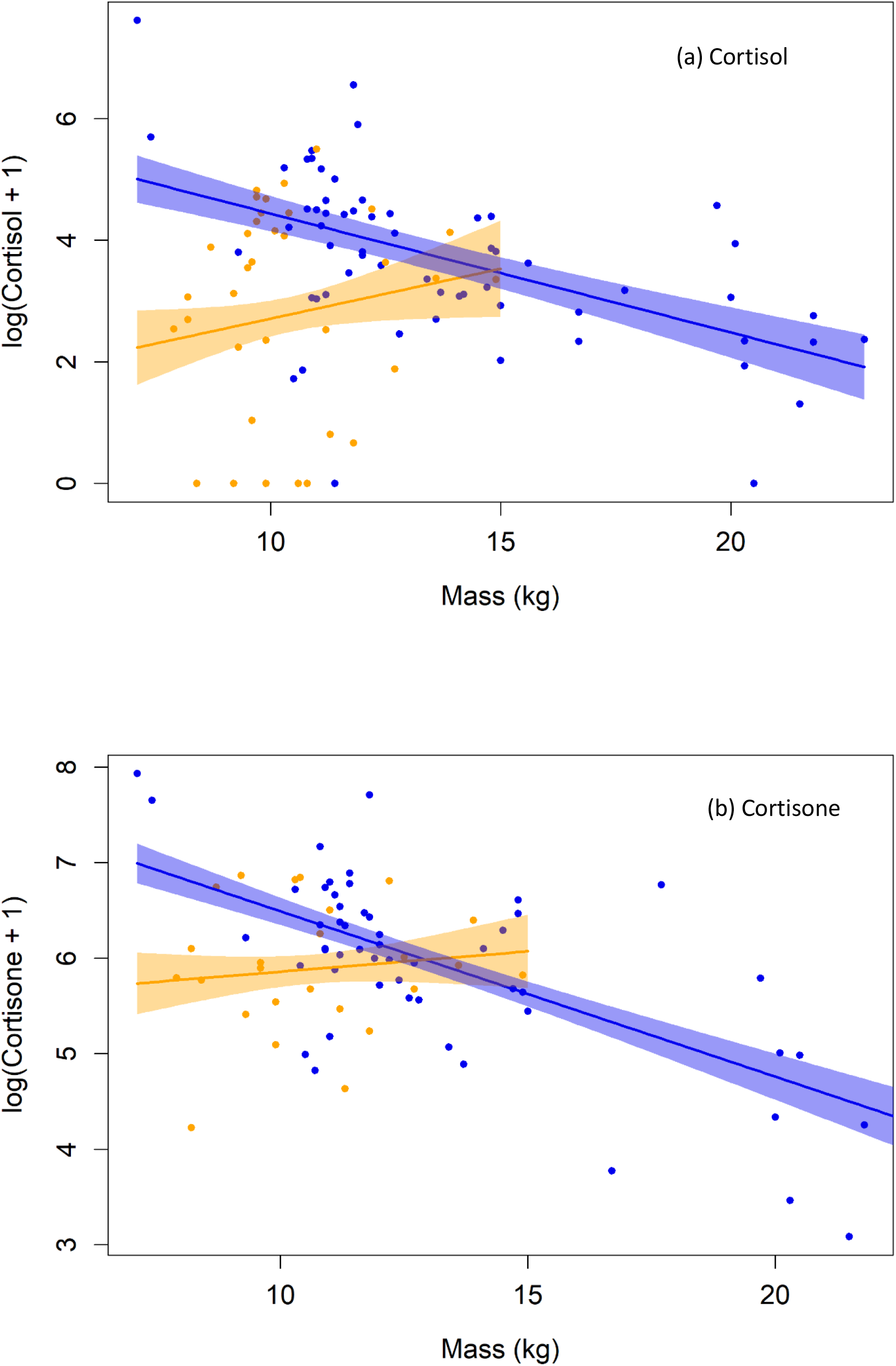
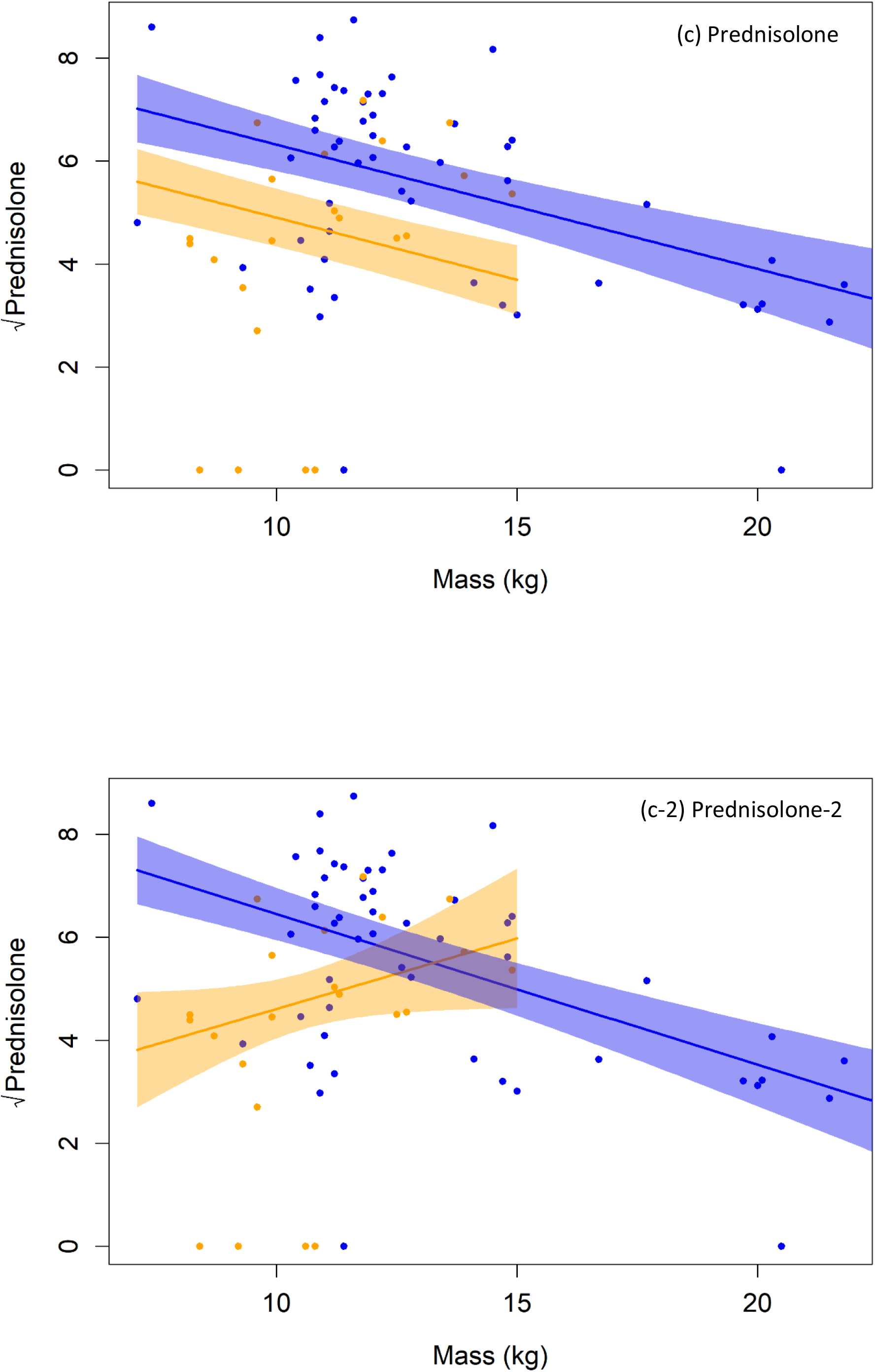
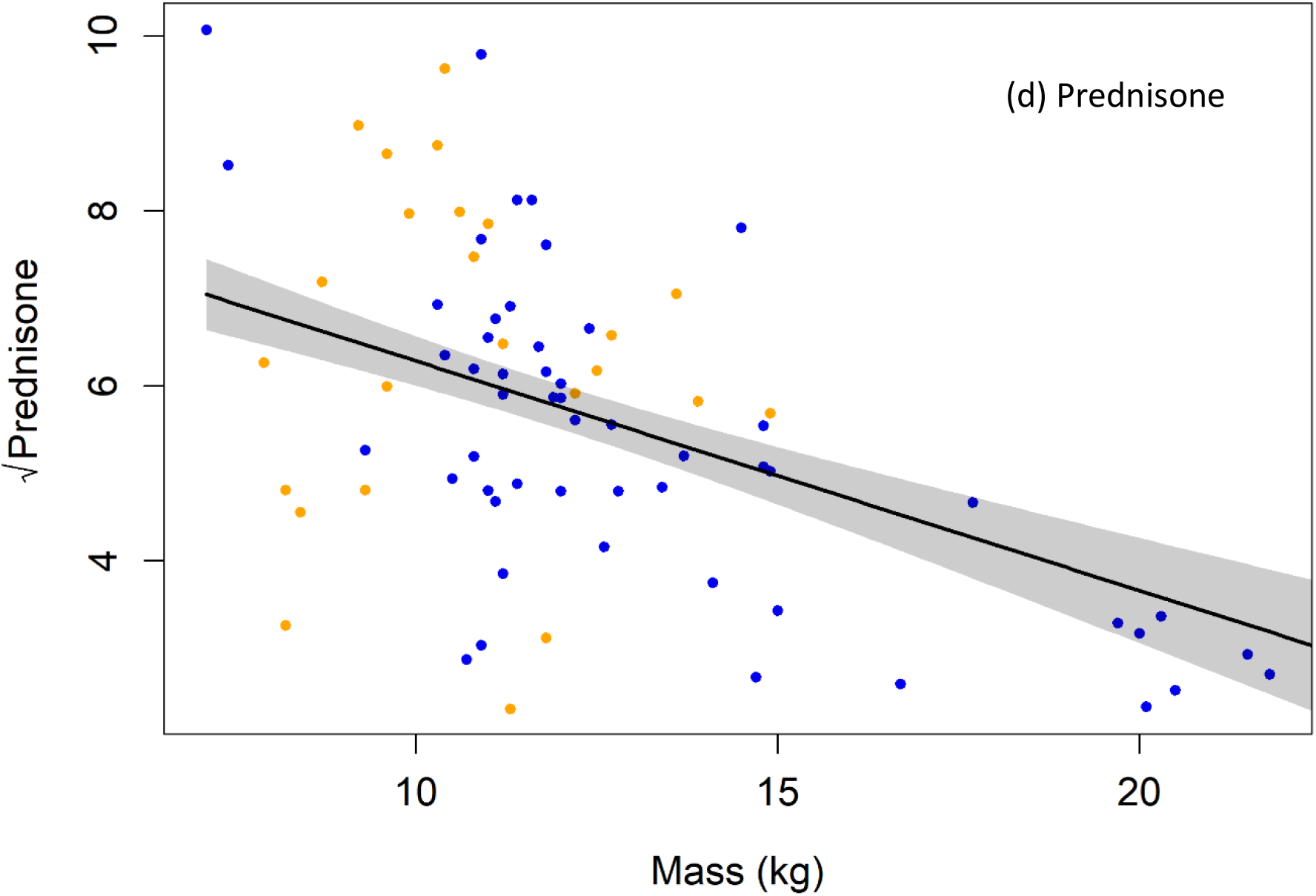
Best-fit linear mixed effects models showing the change in each GC as pups gain mass, from pups provided with water (dark grey) and without water (light grey). Points are GC concentrations (ng/mg Cr), lines are fitted trends, and shaded areas show standard error.

**Table 4.** Results of best-fit linear mixed models explaining the change in GC concentrations in pup urine.

|  | PUP MASS & WATER ACCESS |  |  |  | REHAB DAY & WATER ACCESS |  |  |
| --- | --- | --- | --- | --- | --- | --- | --- |
| BEST-FIT MODELS | % variance | SE | P-value |  | % variance | SE | P value |
| **Log(Cortisol + 1)** | **42.8%** |  |  |  | **42.3%** |  |  |
| Water Access + | 20.0 | 1.85 | **0.006** | Water access + | 21.8 | 0.53 | **<0.001** |
| Pup mass + |  | 0.16 | 0.313 | Rehab day + |  | 0.03 | 0.234 |
| Interaction |  | 0.17 | **0.037** | Interaction |  | 0.03 | **0.005** |
| Pup ID | 22.7 |  |  | Pup ID | 20.5 |  |  |
| **Log(Cortisone + 1)** | **45.4%** |  |  |  | **43.0%** |  |  |
| Water Access + | 38.7 | 0.96 | **0.006** | Water access + | 40.3 | 0.27 | **0.003** |
| Pup Mass + |  | 0.08 | 0.601 | Rehab day + |  | 0.01 | 0.648 |
| Interaction |  | 0.09 | **0.016** | Interaction |  | 0.02 | **<0.001** |
| Pup ID | 6.7 |  |  | Pup ID | 2.8 |  |  |
| **√(Prednisolone)** | **48.7%** |  |  |  | **49.4%** |  |  |
| Water Access + | 8.7 | 0.73 | 0.059 | Water access + | 19.9 | 0.92 | **0.015** |
| Pup Mass |  | 0.09 | **0.007** | Rehab day + |  | 0.04 | 0.490 |
|  |  |  |  | Interaction |  | 0.05 | **0.026** |
| Pup ID | 40.0 |  |  | Pup ID | 29.6 |  |  |
| **√(Prednisolone-2)** | **50.7%** |  |  |  |  |  |  |
| Water Access + | 13.9 | 3.24 | **0.025** |  |  |  |  |
| Pup Mass + |  | 0.28 | 0.334 |  |  |  |  |
| Interaction |  | 0.29 | 0.061 |  |  |  |  |
| Pup ID | 36.9 |  |  |  |  |  |  |
| **√(Prednisone)** | **43.9%** |  |  |  | **43.3%** |  |  |
|  | 24.6 |  |  | Water access + | 34.2 | 0.68 | 0.094 |
| Pup Mass |  | 0.07 | **<0.001** | Rehab day + |  | 0.03 | 0.272 |
|  |  |  |  | Interaction |  | 0.04 | **0.001** |
| Pup ID | 19.3 |  |  | Pup ID | 9.1 |  |  |

Although this interaction was not significant, the second best-fit model explained more variance (R^2^ = 0.507) than the first model (R^2^ =0.487), the fixed effects (pup mass and water access) in the second model explained more variance (13.9%) than the first (8.7%), and the results of the second model were more consistent with that of CL and CN (Table 4; Fig. 7c-2). The predictor variable that best explained PN concentrations was pup mass alone, as PN concentrations decreased significantly as pups gained mass (Table 4, Fig. 7d). The results were similar when rehab day was used in the linear mixed effects models in place of pup mass, but these models explained less variance (Table 4).

## 4. DISCUSSION

### 4.1 The occurrence of prednisolone and prednisone

Our finding of endogenous prednisolone and prednisone in substantial urinary concentrations in rehab harbour seal pups was unexpected. Endogenous urinary prednisolone has been found to occur only in very low concentrations in cattle, with the CL:PL ratio of 25:1 in adult cows (Chiesa et al., 2017), 29–64:1 in veal calves, Famele et al., 2015), and only ~100:1 in humans (Fidan et al., 2013). Endogenous urinary PN has so far been reported as absent in adult cattle (Chiesa et al. 2017, Vincenti et al. 2012), although present in trace amounts (CL:PN ratio 43:1) in a small proportion of veal calves (Famele et al. 2015). The CL:PL and CL:PN ratios close to parity in our rehab seal pups appears to be the first such finding in a mammal species.

### 4.2 Associations between pup urinary GC concentrations and pup growth and body mass

There appeared to be a relationship between GC levels and pup growth and mass, although this was not significant for all GCs. Relatively high prednisolone concentrations were associated with slower growth rates, while prednisone concentrations were higher for pups with mass <11 kg, i.e. below the average healthy birth rate for harbour seals. Moreover, the best-fitting linear mixed effect models indicated that prednisone concentrations were best explained by increasing pup mass alone, while the other GCs showed a non-significant decline with increasing pup mass. This result is consistent with another study, which reported a non-significant association between serum cortisol and increasing body mass (Trumble et al., 2013).

Combining all these results on pup mass and urinary GCs, it seems likely that there may be an effect of slow pup growth and low pup mass on elevated GC prednisolone and prednisone concentrations respectively. The normal wild harbour seal nursing pup growth rate is between 0.4–0.6 kg/day (Skinner, 2006; Cottrell et al., 2002; Muelbert and Bowen, 1993). The closest that the rehab orphan pups in our study came to this was the pups in centre II, with an average growth rate of 0.29 kg/day and consequent relatively high body mass during the sampling period (Table 1); this may be a contributing factor to the pups in centre II having the lowest overall concentrations of prednisolone and prednisone. Nevertheless, the linear mixed effects models indicated that the predictor variables influenced only about 40–50% of prednisolone and prednisone concentrations, and therefore other factors which we were not able to measure may also have been involved.

It is possible that the relatively high levels of endogenous prednisone and prednisolone in the rehab seal pups compared to cattle and humans may be a physiological feature related to the high growth rate in the short lactation period of healthy wild nursing seal pups compared to most land mammals and humans. We hypothesise that when this normal growth rate does not happen and pups remain for a long period with low body mass, that prednisone and possibly prednisolone levels increase to stimulate compensatory glucogenesis. Our results seem to show that prednisone levels fall when pups attain body mass well above healthy birth weight, i.e. when the pups have acquired a significant blubber layer.

We were unable to detect any effect of pup sex on GC concentrations, similarly to the finding of Trumble et al. (2013) with serum cortisol in rehab pups. This is likely because pup mass had a greater influence on GC concentrations, and sexual dimorphism in size is minimal in harbour seal pups (Bowen et al., 1994).

### 4.3 Associations between pup urinary GC concentrations and the rehab physical environment

The outcomes of our linear mixed effects models indicated that urinary cortisol, cortisone and prednisolone (but not prednisone) were negatively associated with the pups having free water access with days in rehab and as they gained body mass. Interestingly, Skinner (2006), studying free-living nursing pups, found that mass gain rate was positively associated with activity in the water. However, in our captive pup study, the interaction between rehab day and having water access was more significant than for body mass and water access.

Although it is not possible in our dataset to clearly distinguish between the effects of body mass and rehab day, it seems likely that as pups become both older and bigger, this in some way creates an increased physiological demand for access to water. If this demand is not met - i.e. pups continue to be maintained in a dry pen – GC levels, particularly of cortisol, may rise. Increasing serum cortisol levels were also recorded in a 1998–2002 group of rehab pups during their first eight weeks in rehab while they were being tube-fed (Trumble et al., 2013). It seems likely that these pups were being maintained without water during this period, although this is not explicitly stated.

Free-living harbour seal pups enter the water less than an hour post-partum (Wilson and Jones, personal observation) and from then on may spend much of their time swimming and diving with their mothers (e.g. Venables and Venables, 1955; Wilson, 1974; Bekkby and Bjørge, 2000), the proportion of their time quantified as 61% by Skinner (2006). This strongly suggests that the physiology of the precocial harbour seal neonate is adapted to an amphibious existence, involving prolonged immersion in sea water (which is around 14°–15°C in July along the Irish and southern North Sea coasts; https://www.seatemperature.org). It is therefore likely that young harbour seal pups’ physiology may become disordered when they are maintained without water access. There is some evidence that the normal post-natal development of red blood cells increasing in size to enable storage of enough haemoglobin, and hence oxygen to fuel diving, is dependent on the pup “practicing” diving and apnoea during the early post-natal period (Thompson and Ono, 2015). These authors suggest that pups undergoing rehab ideally, therefore, require water of sufficient depth for them to submerge for extended periods, and indicated that the orphan pups at their rehab centre normally acquire this experience as they play, socialise in the water - and sometimes sleep while submerged. These conditions were provided provided in centres II and IV. The centre II pups, with a choice of a shallow paddling pool or deeper bath, invariably chose the bath to play (Fig. 2) and (as they grew older) to sleep while submerged (SW, personal observation).

### 4.4 The relevance of this study to the question of cortisol levels and welfare

The normal behaviour of free-living harbour seal pups with their mothers involves continual mutual tactile and olfactory contact, principally in the water or at the water’s edge (e.g. Wilson and Kleiman, 1974; Wilson & Jones, 2018). It may therefore be a reasonable assumption that orphan pups maintained with one or more partner pups and free water access allows for the expression of natural filial and motor behaviours to the greatest extent possible, and therefore maximal welfare within the inevitable confines of a rehab environment. Conversely, orphan pups maintained in isolation and without water access may, according to the same benchmark, be considered as poor welfare. Our initial hypothesis (in 2012–13) was therefore that isolated/dry-pen pups would have higher urinary cortisol levels than the social/free-water-access pups. The initial results for cortisol alone, comparing the 2012–2013 results from centres I and II, seemed to support this. However, further analysis of samples that included three further centres taken between 2014–2016 indicated that the cortisol-welfare relationship regarding these orphan seal pups was more complex.

Free-living pups losing contact with their mothers in the water typically give the distress call (Perry and Renouf, 1988; Di Poi et al. 2015) accompanied by agitated swimming as they try to reconnect with their mother. It has been shown that free-living nursing harbour seal pups separated briefly from their mothers had elevated plasma cortisol levels (Di Poi et al., 2015). This is consistent with experimental results with separating guinea-pig infants and is called the “acute” phase of maternal separation (Yusko et al., 2012).

However, in our study we were taking samples from maternally deprived pups held in isolation for extended periods in centres I and III, and from pups - also maternally deprived, but provided with companion pups and free water access in centres II and IV. The isolate/dry-pen pups in centre III had the lowest average concentrations of urinary cortisol of all the centres, whereas the social-water-access pups of centre IV had the highest levels (Figure 4). This immediately indicates that our original hypothesis was too simplistic.

The maternally deprived pups held in isolation would be expected to be in the post-separation “depressive” phase, as in rhesus monkeys (Spencer-Booth and Hinde, 1971), when the separated infants show reduced locomotion and other passive behaviours. Reduced cortisol and adrenocortical activity have been shown in captive infant primates reared without their mother (Novak et al., 2013) and cortisol levels have been shown to fall in guinea-pig infants during the depressive phase following maternal separation (Yusko et al. 2012). Feedback mechanisms may result in reduced cortisol output in chronic or sustained stress situations (Sapolsky et al., 2000; Novak et al. 2013), including in harbour seal pups in rehabilitation (Gulland and Haulena, 1999). Chronic conditions in adult animals, caused by repeated or sustained stress, are a state of “distress”, which may be manifest in depressed cortisol levels (Linklater et al. 2010). Such complexities involving stress, depressive phase, distress and feedback may have contributed to the variable cortisol levels found in pups in centres I and III, all of which were held in isolation in dry pens.

Orphan pups in captivity housed with one or more other pups and water access appear to act as substitute mothers to one another, (S.Wilson and R. Alger, unpublished data), thereby preventing the depressive phase of separation and reducing or eliminating distress. Although most of the pups in our study with free water access were kept socially, some pups were maintained socially but without water (Fig. 1, Table 1), which might explain, at least partially, why water access but not social group appeared to influence GC levels, particularly of cortisol and cortisone.

Since free water access was almost always accompanied by social housing in this study, it seems likely that the combination of both factors (in centres II and IV) – or the absence of both (centres I and III) may have influenced the cortisol and cortisone levels observed. The paired pups in centre II all had a stable pair-bonded relationship, which was not observed in centre IV pups living in groups (personal observation). The vigorous social play in the water observed by the pups of this study in centre II may be fuelled partly by glucogenesis due to adrenal stimulation, thereby maintaining a measurable level of cortisol production. It is also possible that the constant social contact involved in the pair bond of the centre II pups may have a “stress buffering” role (Morrison 2016). The extremely low cortisol levels in the isolated/dry-pen pups in centre III might suggest chronic stress or distress in these isolated/dry-pen pups, although the similarly maintained centre I pups had relatively elevated cortisol levels. All these possible factors in combination complicates a welfare interpretation of cortisol levels.

The continual contact of the pair-bonded pups, with presumed oxytocin release, may have contributed to the relatively rapid mass gain in centre II pups (Robinson et al., 2019). This early mass gain allowed for a relatively early release, thus facilitating natural behaviour patterns of foraging and social haul-out following release (Wilson, 1999; Greig et al., 2019).

The reasons for the relatively elevated cortisol levels in centre IV compared with centre II (Fig. 4) are not obvious in the context of the aspects of the social and physical environment already considered, since both sets of pups were provided with free water access and most centre IV pups were also kept socially. A possible partial explanation of the relatively high levels of cortisol, cortisone and prednisolone in centre IV pups might involve elevated levels of persistent organic pollutants (POPs) recorded in the Wadden Sea. (Laane et al., 2013; Weirup et al., 2013). POP contaminants in maternal tissues may be transferred to the foetus, posing health risks during development (Wang et al. 2012).

## 5. CONCLUSIONS

This is the first study to investigate the concentrations of four GCs in urine of harbour seal pups held in rehabilitation centres. We found that different stressors may affect the differential release of the four GCs: prednisone appeared to be most strongly implicated in relation to low body mass, whereas increasing levels of the other three GCs were associated with both body mass and lack of free water access. Our results indicate the potential utility of individual GC analysis in assessing rehab pups’ physiological welfare, and highlight the importance of water access to harbour seal pups in rehabilitation. Further research is required to investigate GC concentrations in orphans of harbour seals and other phocid species to reveal a more complete picture of the role of GC concentrations in the welfare of orphans in rehabilitation centres.

## 6. ACKNOWLEDGEMENTS

We are grateful to Joanna Keenan for carrying out the initial UPLC-MS/MS analyses at Queens University, N. Ireland, and to Brett Greer for his help in facilitating the analyses. We would especially like to thank Professor Chris Elliott at the Institute for Global Food Research, Queens University, and Dr Robert McCracken at the Agri-Food & Biosciences Institute, N. Ireland, for permitting the analyses in their departmental laboratories.

We are indebted to all the personnel at the seal sanctuaries participating in this study for their generosity in permitting and facilitating the urine sampling. In particular we would like to acknowledge the invaluable support of Duncan, Richard and Nicki Yeadon and Curtis Jones at Natureland, Skegness, England; Will Matthews during the study period at Skegness, Fleur Brochut at Tara Seal Research, N. Ireland; Tania Singleton at Exploris, N. Ireland; Tanja Rosenberger and Janne Sundermeyer at Seal Centre Friedrichskoog, N. Germany, and Ally McMillan, Lisa Becker and Sonja Ciccaglione at Seal Rescue Ireland, Wexford, Ireland.

## ETHICS STATEMENT

The urine sampling in this study was carried out with the permission and collaboration of each of the five seal centres. Four of the five centres (I, III, IV and V) were in the public domain and acting within relevant government guidelines. Centre II was a voluntary private centre operating with local government (N. Ireland) permission. The sampling did not cause additional disturbance to the pups beyond their routine care and handling and did not impact their welfare. No specific license for this study was therefore required.

