## Supplementary file S1 for "Urinary glucocorticoids in harbour seal (*Phoca vitulina*) pups during rehabilitation"

SUPPORTING FILE S1

METHOD AND METHOD VALIDATION FOR GLUCOCORTICOID STEROID DETERMINATION BY ULTRA-PERFORMANCE LIQUID CHROMATOGRAPHY TANDEM MASS SPECTROMETRY (UPLC–MS/MS) IN TWO LABORATORIES PARTICIPATING IN THE STUDY OF HARBOUR SEAL PUP URINE

1. **Institute of Global Food Research, Queens University, Belfast (QUB) (Laboratory 1 in main text)**

***UPLC-MS/MS quantification of cortisol (CL) in urine***

Laboratory 2 quantified the cortisol content of 11 duplicate urine samples from Centre I (2012) and a further 8 samples from Centre II (2013) (see main text s. 2.2–2.3 for explanation of Centres and sampling) by ultra-performance liquid chromatography tandem mass spectrometry (UPLC–MS/MS).

This method is transcribed here from Keenan (2015) and describes the UPLC–MS/MS procedure and method validation for quantifying cortisol (CL) in bovine urine. The analysis of the seal pup urine for CL in the present study (Lab 2 did not analyse the seal pup samples for cortisone) was carried out alongside the ongoing bovine urine analyses in that separate study. The parallel procedures for quantifying cortisone are therefore omitted from the following description.

*Measurement of the urinary creatinine (Cr) concentration of the samples*

Creatinine (Cr) is commonly used as a correction factor to control for different urinary volumes as it is eliminated through urine at a constant rate and is reflective of excretion rates (Novak et al. 2013). Urinary GC concentrations are expressed as a ratio to the creatinine concentration (ng GC/mg Cr) in each sample. Creatinine concentrations in each sample were analysed using a colorimentric method modified from the Jaffe reaction, in which Cr in an alkaline solution reacts with picric acid to form a coloured complex (Taussky and Kurrzmann 1954).

Three assay reagent mixtures were produced to perform creatinine analysis: Solution A was a mixture containing sodium hydroxide (final concentration 0.275 M), dibasic sodium phosphate (final concentration 0.06 M) and disodium tetraborate (final concentration 0.06 M). Solution B consisted of sodium dodecyl sulphate (final concentration 0.1 M) and Solution C consisted of picric acid (final concentration 0.06 M). A 1 mg ml^-1^ stock Cr standard was prepared in 0.1 M hydrochloric acid (HCl). A dilution schedule was followed to produce working standards of 500 µg ml^-1^ to 0 µg ml^-1^ in 0.1 M HCl. A 10 µl volume of urine and standard were added to wells in a 96 well plate. Equal volumes of reagents A, B and C were added together at the point of analysis and 200 µl of this mixture added to each well. After reaction time of 30 min, absorbance was read on a spectrophotometer at 510 nm. Following this, 25 µl of 30% acetic acid were added to each well and left at room temperature for 5 min and a final absorbance taken. The second absorbance was subtracted from the first absorbance to determine urinary Cr concentrations, which were quantified using a linear calibration regression curve in the range 0 µg/ml – 500 µg/ml.

*Reagents and Materials* *for UPLC-MS-MS*

Analytical grade reagents and HPLC grade solvents were used throughout. HPLC grade methanol, ethyl acetate, acetonitrile and formic acid were purchased from Fisher Scientific (Loughborough, UK). HPLC grade hexane, 25% ammonium hydroxide, acetic acid, activated charcoal and analytical standards (≥98% HPLC hydrocortisone and the deuterated standard d4-hydrocortisone were all purchased from Sigma Aldrich (Dorset, UK). LC-MS grade water (18.2Ω) was produced from a Milli-Q Millipore water system.

*Sample preparation and extraction of urine for UPLC-MS/MS*

Sample preparation and UPLC-MS/MS conditions were modified from a previous method described by McWhinney et al. (2010). Prior to analysis, urine was defrosted at room temperature, 1ml placed into Eppendorf vials, and centrifuged at 10,000 x g for 10 min at 4°C. 100 µl of centrifuged urine was then removed and added to 530 µl of LC-MS grade water. In addition, 10 µl of 500 ng/ml d4-hydrocortisone (CL) were added to each sample as internal standards, to produce a total volume of 650 µl for loading on to Oasis HLB 1 cc (30 mg) solid phase extraction (SPE) cartridges (Waters Corporation, Milford, MA, USA). SPE cartridges were activated with 1 ml methanol followed by 1 ml water and urine samples loaded and washed sequentially with 1 ml 20% methanol, 1 ml hexane and eluted with 2 x 1 ml ethyl acetate. Subsequently, the elute was dried down under nitrogen at 35°C for 15 min. using a TurboVap LV (Caliper Life Sciences, Hopkinton, USA). All dried extracts were reconstituted with 100 µl of 45% methanol containing 2mM ammonium acetate and 0.1% (v/v) formic acid and transferred to LC vials prior to analysis by mass spectrometry.

*Urinary extracted matrix-matched calibration curve*

Urine extracted matrix-matched curves were prepared using steroid-free urine for the purposes of CL quantification. Similar to real urine samples, the steroid-free urine was centrifuged at 10,000 x g for 10 min at 4°C. 100 µl of urine was added to 430 µl of water and 10 µl of 500 ng/ml of each deuterated standard. 50 µl of CL standards were added to charcoal-stripped urine and LC-MS water to create a total of seven calibrants ranging from 3.9 ng/ml to 250 ng/ml. Calibrants were extracted using the afore-mentioned extraction (SPE) process. The calibration curve was plotted using the response factor: analyte area * (internal standard concentration / internal standard area).

To assess the effectiveness of the charcoal stripping process, a quality control sample was spiked with 10 µl of d4-CL and included in every mass spectrometry run. During the quantification process this sample was included as a standard in the extracted matrix-matched calibration curve as a concentration of 0 ng ml^-1^ so that it could be quantified against itself. Following this, the charcoal-stripped sample was excluded from the calibration curve and the quantified value added to the nominal concentration of all standards (standard addition), hence correcting for the analyte level that had not been completely stripped in the calibration curve.

*UPLC-MS/MS analysis of bovine urine (the same method was used for seal pup urine)*

UPLC-MS/MS analysis was performed on an Acquity UPLC system with a binary solvent manager that was coupled to a Micromass Quattro Premier XE mass spectrometer (Waters Corporation, Milford, MA, USA).

10 µl of urine extract was injected on to an Acquity BEH C18 column (1.7 µm, 2.1 mm x 50 mm) equipped with a pre-filter. Column temperature was maintained at 50°C and autosampler temperature was maintained at 8°C. Separation was achieved at a flow-rate or 0.4 ml/min with mobile phase A consisting of water with 2 mM ammonium acetate and 0.1% (v/v) formic acid and mobile phase B consisting of methanol with 2 mM ammonium acetate and 0.1% formic acid. The solvent gradient used is shown in Table 1.1.

Table 1.1. Mobile phase A and B gradient conditions for cortisol and cortisone separation in bovine urine or blood using UPLC-MS/MS.

| **Time (min)** | **%A** | **%B** | **Curve** |
| --- | --- | --- | --- |
| Initial | 55.0 | 45 | Initial |
| 0.50 | 55.0 | 45.0 | 6 |
| 2.00 | 50.0 | 50.0 | 6 |
| 2.50 | 5.0 | 95.0 | 6 |
| 3.00 | 5.0 | 95.0 | 6 |
| 3.10 | 55.0 | 45.0 | 6 |
| 4.00 | 55.0 | 45.0 | 6 |

The mass spectrometer was maintained in electrospray ionisation positive mode (ESI+) with multiple reaction monitoring (MRM) acquisition, the transitions of which are illustrated in Table 2. The capillary voltage was set at 2.50 kV and cone voltage at 30 V. Flow rates of the desolvation gas and cone gas was 700 L/h and 50 L/h respectively. The source temperature was maintained at 120°C and desolvation temperature at 350°C. All data were processed using MassLynx V4.1 (TargetLynx) software (Waters Corporation, Milford, MA, USA).

Table 1.2. MS/MS parameters utilised for the targeted analysis of cortisol and cortisone in bovine urine or blood

| **Compound** | **Transition** | **Cone voltage (V)** | **Collision energy (eV)** |
| --- | --- | --- | --- |
| Cortisol (quantifier) | 363.25>120.90 | 30.0 | 20.0 |
| Cortisol (qualifier) | 363.25>309.20 | 30.0 | 18.0 |
| d4-Cortisol | 367.20>121.00 | 30.0 | 25.0 |
| Cortisone (quantifier) | 361.10>120.95 | 30.0 | 35.0 |
| Cortisone (qualifier) | 361.10>163.00 | 30.0 | 25.0 |
| d2-Cortisone | 363.10>165.05 | 35.0 | 25.0 |

*UPLC-MS/MS method validation*

UPLC-MS/MS method validation was undertaken and accuracy/recovery, intra- and inter-day repeatability and linearity were assessed. Charcoal-depleted steroid free urine was spiked with 30 µl, 20 µl and 4 µl of a 250 ng/ml CL and CN solution to produce levels of 200 ng/ml, 50 ng/ml and 10 mg/ml respectively. Six replicates of all three concentrations were extracted using the previously mentioned solid phase extraction (SPE) process and analysed by UPLC-MS/MS to determine within run precision (intra-day repeatability). This was repeated on a further two days to assess day to day precision (inter-day repeatability).

Linearity of the response was assessed by extracting matrix-matched calibration curves, with a total of seven calibration points ranging from 3.9 ng/ml to 250 ng/ml for both CL and CN. The regression coefficients (R^2^) for both calibration curves used within this study were >0.998, demonstrating excellent linearity.

Although not necessary for natural hormones, method validation was applied as outlined in accordance with EU guidelines (2002/657/EC) so that the decision limit (CCα) and detection capability (CCβ) could be employed. The decision limit (CCα) was determined by using the intercept of the calibration curve procedure. This involves the value of y (response factor), where x (concentration) is equal to zero and 2.33 times higher than the standard error of intercept for a set of data that has six replicates of three different concentrations. Calculation of the detection capability (CCβ) involved the addition of 1.64 times the standard error to the decision limit (CCα) (European Commission, 2002).

*References*

European Commission (2002) 2002/657/EC: Commission Decision of 12 August 2002 implementing Council Directive 96/23/EC concerning the performance of analytical methods and the interpretation of results (Text with EEA relevance) (notified under document number C(2002) 3044). Available from: <https://publications.europa.eu/en/publication-detail/-/publication/ed928116-a955-4a84-b10a-cf7a82bad858/language-en>

Keenan JB (2015) Metabolomic approaches to assessing pre-slaughter stress in relation to beef quality. PhD Thesis, University of Reading, UK. Available from: uk.bl.ethos.679250.

McWhinney BC, Briscoe SE, Ungerer JPJ, Pretorius CJ (2010) Measurement of cortisol, cortisone, prednisolone, dexamethasone and 11-deoxycortisol with ultra high performance liquid chromatography–tandem mass spectrometry: application for plasma, plasma ultrafiltrate, urine and saliva in a routine laboratory. J Chromatogr B. 2010; 878: 2863–2869.

Novak, MA, Hamel, AF, Kelly BJ, Dettmer AM and Meyer JS. Stress, the HPA axis, and non-human primate well-being: a review. Appl Anim Behav Sci. 2013; 143: 135-149.

2. **Chemical and Immunodiagnostic Sciences Branch, Veterinary Sciences Division, Agri-Food & Biosciences Institute (AFBI), Belfast (Laboratory 2 in main text)**

***UPLC-MS/MS quantification of cortisol (CL) in urine***

Laboratory 2 analysed 98 samples collected from centres II, III, IV and V in 2014–16 for urinary concentrations of cortisol (CL), cortisone (CN), prednisolone (PL) and prednisone (PN) by ultra-performance liquid chromatography tandem mass spectrometry (UPLC/MS/MS), according to the Laboratory 2 analysis of bovine liver as part of the routine statutory Veterinary Drug Residue testing programme at AFBI, in accordance with EU guidelines (2002/657/EC). The seal pup urine samples were analysed according to the same procedures used in the statutory drug testing system.

*Measurement of the urinary creatinine (Cr) concentration of the samples*

The creatinine in the 98 samples from Centres II–V collected in 2014–16 were analysed using an Audit Diagnostics Creatinine Jaffe Reaction kit.

The samples were tested by an automated system on an Audit Diagnostics Sapphire 800 analyser. The automated sampler takes up 15 µl of undiluted urine and transfers it to a reaction cell. 250 µl of reagent 1 (Alkaline buffer) is added and mixed. 50 µl of reagent 2 (picric acid) is added and mixed. The reaction time is 10 minutes and measurements recorded ‘2-point rate’ with main wavelength 505 nm and second wavelength 570 nm.

*Quality control.* The analyser is calibrated daily using the Audit Diagnostics General Chemistry calibrator. Calibration is 2-point linearity. The analyser performance is quality controlled using the Audit Diagnostics General Chemistry level 1 (low) and level 2 (high) QC controls.  These have assigned values based on each lot number of the kit used. The QC values are taken each day and recorded in the lab database. For an analysis to be valid, QC must fall within 1.0 standard deviation of the expected value.

*Reagents and Materials* *for UPLC-MS-MS*

Analytical grade reagents and HPLC grade solvents were used throughout. Methanol was obtained from Romil (Cambridge UK), acetonitrile, propan-2-ol and sodium hydroxide (NaOH) pellets from Fisher Scientific (Leicestershire, UK), 1M HCl from BDH (Dorset, UK), and ammonium acetate from Sigma Aldrich (Dorset, UK). Grade 1 (18MΏ.cm) water was obtained from an in-house Milli-Q system (Millipore corp., Livingston, UK). Solid-phase extraction was performed on Oasis HLB SPE columns (Waters Chromatography Ireland Limited). Screw capped polypropylene centrifuge tubes were used for hydrolysis. Analytical standard powders for CL, CN, PL and PN were obtained from Sigma-Aldrich. Isotopically labeled internal standard powders for d6-prednisolone and d4-cortisol were obtained from CDN isotopes (Quebec, Canada).

Individual stock standard solutions were prepared at 1 mg/ml in methanol. Mixed working standard solutions for CL, CN, PL and PN were prepared at 1µg/ml and 100 ng/ml by dilution of the stock standard with methanol, and a mixed internal standard spiking solution for d6-PL and d4-CL at 1 µg/ ml in methanol. Standards were stored in 25ml amber glass vials at 4ºC for up to 12 months. The mixed 100 ng/ml and 1 µg/ml working standards were used to prepare matrix matched calibration curves across the range 0.5–10.0 µg/kg (ng/ml equivalent).

Additional reagent solutions were:

- 0.2 M NaOH (prepared fresh weekly and stored at room temperature)
- 0.02 M NaOH/methanol (60/40 v/v) and 5mmol ammonium acetate solution in methanol (prepared fresh weekly and refrigerated)
- UPLC mobile phase A: 0.05 mM ammonium acetate in 10% methanol (prepared fresh daily)
- UPLC mobile phase B: acetonitrile (fresh for each analysis)
- UPLC needle wash: methanol (30%), acetonitrile (30%), propan-2-ol (30%) and water (10%) (prepared fresh for each analysis).

*Sample preparation and extraction of urine for UPLC-MS/MS*

Bovine urine was used as the control negative, and for the matrix standard curve. On the day of analysis the seal urine samples and control negative were defrosted at room temperature and mixed thoroughly prior to dispensing of aliquots.

Seal urine (5 ml) was pipetted into a 50 ml screw-cap polypropylene centrifuge tube, while bovine urine samples were set up for negative and positive controls and for matrix standards. All seal urine, and all positive/negative controls were spiked with 20 µl of the 1 µg/ml mixed internal standard. The selected bovine urine blank samples for recovery evaluation were fortified with 100 µl of the mixed 100 ng/ml standard and allowed to equilibrate for 10 minutes before proceeding.

Methanol (10ml) and 1M Hydrochloric acid (15 ml) was added to each tube, which was then capped and vortexed for 20 sec. All tubes were incubated in a water bath for 4 hours at 50ºC. After incubation tubes were allowed to cool to room temperature and centrifuged at 3500 rpm for 15 min.

Solid phase extraction (SPE) was performed using Oasis HLB 3cc, 60 mg SPE cartridges, placed on a Vac-Elute manifold. Each cartridge was conditioned with methanol (3 ml) followed by water (3 ml) (cartridges were not allowed to dry out at any stage until the final wash). The sample extract was applied and allowed to pass through under gravity. The cartridges were washed with water (3 ml) and allowed to dry by passing air through under vacuum for 5 min. The samples were eluted into 13*100 mm disposable glass tubes with methanol (3 ml). At this stage the matrix standard tubes were spiked with the mixed 100 ng/ml and 1 µg/ml standards, and with the 1µg/ml mixed standard (Table 2.1).

Table 2.1. Volumes of standard solution used to spike the matrix standard tubes at each concentration

| Standard concentration (ng/ml equivalent) | Volume (µl) of 100 ng/ml mixed quaternary std | Volume (µl) of 1 µg/ml mixed tertiary std | Volume (µl) of 1 µg/ml mixed internal tertiary std |
| --- | --- | --- | --- |
| 0.5 | 25 | - | 20 |
| 1.0 | 50 | - | 20 |
| 2.0 | 100 | - | 20 |
| 5.0 | - | 25 | 20 |
| 10.0 | - | 50 | 20 |

All sample, control and matrix standard tubes were evaporated on a TurboVap, under nitrogen, at 55 ºC, and reconstituted in methanol (250 µl), vortexed for 20 seconds, followed by water (250 µl). Extracts were transferred to a 2 ml vial with tapered insert prior to UPLC-MS/MS analysis.

*Instrumentation and conditions.* UPLC separations were carried out using an Agilent 1290 infinity LC system comprising a G4220A Binary Pump, G1330B Thermostat, G4226A Sampler, and a G1316C Thermo-stated Column Compartment (Agilent Technologies, Inc. Santa Clara, USA). A reverse phase gradient separation was achieved on an Agilent EclipsePlusC18, RRHD 1.8 µm, 2.1*100mm chromatographic column (Agilent Technologies, Inc. Santa Clara, USA) with an in-line filter assembly (0.5 µm porosity) (Waters, Milford, USA). The sample compartment was set at 10ºC, column temperature at 40ºC, and an injection volume of 1 µl used. The mobile phases were (A) 0.05 mM ammonium acetate in 10% methanol and (B) acetonitrile, with a flow rate of 0.4 ml/min. The flow was diverted to waste during sample injection cycles at 0.0 – 2.5 min and again from 9.4 min until the end of an injection cycle. Linear gradient steps were used with initial conditions set at 80% A, decreasing to 68.5% A after 2 min, with a 1 min hold at 68.5% A, and returning to 80% A at 9.0 min. A re-equilibration period of 2.6 min was used. Total analysis time was 11.6 min per sample.

An Agilent AG6490 triple-quadrupole controlled by Mass-Hunter acquisition software was used for mass spectrometric analysis; this was connected to the UPLC system via an electrospray ionisation (ESI) interface source. All compounds were analysed in negative ESI mode with Jet Stream. The following source parameter conditions were used: Gas Temp 200 ºC, Gas Flow 12 L/min, Nebulizer 50 psi, Sheath Gas 350ºC, and Capillary and Nozzle voltages 4000V and 2000V respectively in in negative mode. Nitrogen was used for nebulizing, sheath, drying and collision gas. Detection was performed in dynamic multiple reaction monitoring mode (DYN-MRM) at unit mass resolution with a Cycle Time of 300 msec. The MRM experiment is summarised in Table 2.2. The most intense MRM was used as the quantitative ion, with the remaining MRMs being used for ion ratio confirmation.

Table 2.2. Summary of multiple reaction monitoring (MRM) experiment

| **Compound Name** | **Precursor**  **Ion** | **Product**  **Ion** | **Collision**  **Energy (V)** | **Ref. Time (min)** | **Ref.**  **Window** | **Polarity** |
| --- | --- | --- | --- | --- | --- | --- |
| Prednisone | 417.1 | 327 | 8 | 4.23 | 0.59 | Negative |
| Prednisone | 327.1 | 285.1 | 20 | 4.23 | 0.59 | Negative |
| Prednisone | 327.1 | 148.8 | 36 | 4.23 | 0.59 | Negative |
| d6 Prednisolone | 333.1 | 284 | 36 | 4.31 | 0.51 | Negative |
| Prednisolone | 419.1 | 329.1 | 24 | 4.39 | 0.46 | Negative |
| Prednisolone | 328.1 | 295.2 | 20 | 4.39 | 0.46 | Negative |
| Prednisolone | 329.1 | 280 | 36 | 4.39 | 0.46 | Negative |
| Cortisone | 419.1 | 329.1 | 12 | 4.53 | 0.82 | Negative |
| Cortisone | 329.1 | 301.2 | 8 | 4.53 | 0.82 | Negative |
| Cortisone | 329.1 | 136.9 | 28 | 4.53 | 0.82 | Negative |
| d4 Cortisol | 335.1 | 301.2 | 20 | 4.53 | 0.6 | Negative |
| Cortisol | 421.1 | 331.1 | 12 | 4.56 | 0.71 | Negative |
| Cortisol | 421.1 | 297.1 | 36 | 4.56 | 0.71 | Negative |
| Cortisol | 331.1 | 297.2 | 20 | 4.56 | 0.71 | Negative |
| Cortisol | 331.1 | 282.1 | 28 | 4.56 | 0.71 | Negative |

Results were processed using Mass-Hunter quantitative software. Internal standard correction was applied for PN and PL (d6-prednisolone), and CN and CL (d4-cortisol).

*UPLC-MS/MS method validation*

The validation method in Lab 3 is performed as part of the routine statutory Veterinary Drug Residue testing programme at AFBI, for which the designated tissue of choice is liver. For bovine liver validation, three sets of six replicate blended negative liver samples were spiked at 1.0 ppb, 2.0 ppb and 3.0 ppb and extracted and analysed by the above procedure against a matrix spiked calibration curve over the range 0.5 µg/kg –10 µg/kg. Liver samples were homogenised for 30 seconds using a Silverson SL2 laboratory homogeniser (for seal pup urine analysis in the present study, the samples were vortexed).

The extraction was repeated on three separate days, and after ensuring all relevant identification criteria were met, the collated data were used to determine recovery, within-day, between-day and intermediate precision, and - by using the intercept of the calibration curve method - the parameters CCα (decision limit) and CCβ (detection capability). As corticosteroids are not authorised for use in the EU, the alpha-error was set at 1%.  To determine CCα a calibration line is produced based on all the contributing recovery responses at the three levels 1.0 ppb, 2.0 ppb & 3.0 ppb over the three days. This is extrapolated to determine the signal (y response) at concentration x equal to zero; the Standard Error of the intercept is determined and multiplied by 2.33 (1%, one sided) to give a response equivalent to a concentration CCα. The detection capability is equivalent to the concentration obtained from a signal at CCα plus 1.64 times the Within-Laboratory reproducibility at CCα.
