## Supplementary file S2 for "Urinary glucocorticoids in harbour seal (*Phoca vitulina*) pups during rehabilitation"

METHOD FOR CORTISOL DETERMINATION BY ENZYME-LINKED IMMUNOABSORBANT ASSAY (ELISA)

**ELISA assays carried out at the University of Lincoln, Riseholme Park, Lincolnshire, UK**

Enzyme-linked immunoabsorbent assay (ELISA) was used to quantify the cortisol content of 17 urine samples from two male pups in rehabilitation Centre I (2012) (see main text s. 2.2–2.3 for explanation of Centres and sampling). Here we describe the steps by which these ELISA analyses were carried out, and compare the results with those obtained from 11 duplicate samples by UPLC-MS/MS (see main text s. 2.4).

*Measurement of the urinary creatinine (Cr) concentration of the samples*

Creatinine (Cr) is commonly used as a correction factor to control for different urinary volumes as it is eliminated through urine at a constant rate and is reflective of excretion rates (Novak et al. 2013). Urinary GC concentrations are expressed as a ratio to the creatinine concentration (ng GC/mg Cr) in each sample. Creatinine concentrations in each sample were analysed using a colorimetric method modified from the Jaffe reaction, in which Cr in an alkaline solution reacts with picric acid to form a coloured complex (Taussky and Kurzmann 1954).

A standard of 0.1 mM/L Cr solution was prepared to be used as a reference from a stock solution of 10 mM Cr (113.12 mg creatinine dissolved in 100 ml of 15MΩ water). A 750 mM/L sodium hydroxide (NaOH) solution was prepared by dissolving 3g of NaOH in 100 ml of 15MΩ water. The urine samples were first defrosted to room temperature. 1.5 ml of diluted urine sample was added to a 4ml PMMA cuvette, followed by 0.5 ml of 1.3% (saturated) picric acid in water. A standard cuvette was prepared with 1.5 ml of 0.1 mM/L Cr solution in place of urine. A blank cuvette was prepared with 1.5 ml of 15MΩ water in place of Cr. The colour reaction was started by addition of 0.5 ml of 750 mM/L NaOH to each cuvette. The absorbance of each sample was then measured at a wavelength of 550 nm after 25 min using a WPA Biowave II UV-Vis spectrophotometer, that had been zeroed using a cuvette containing only distilled water. The absorbance rise (urine cuvette – blank cuvette) was divided by the absorbance rise (standard cuvette – blank cuvette) and multiplied by 10 to obtain the concentration of creatinine (mM/L) in the undiluted urine.

*Analysis of urinary cortisol (CL) concentrations* *by ELISA*

In an ELISA (enzyme-linked immunoabsorbent assay) for the determination of cortisol (CL) concentration in urine, a specific antibody binds to an enzyme-labelled target antigen (CL in this case). The enzyme activity is measured using a solid substrate that changes colour when modified by the enzyme. The intensity of the colour depends on the CL concentration and is read on a specific type of microplate. Urinary CL concentrations in 17 samples from centre I in 2012 were determined using a cortisol ELISA kit (Cat No ADI-900-071) for colorimetric detection from Enzo Life Sciences (UK) Ltd. The kit is stated to be appropriate for plasma, saliva, serum or urine from any species. The manufacturer this ELISA kit for cortisol (CL) determination is reliable, reproducible, sensitive, and has been used in numerous published and highly cited animal and human studies (<http://www.enzolifesciences.com/ADI-900-071/cortisol-elisa-kit/>). The cross-reactivity with other steroid hormones is stated to be Cortisol (100%), Prednisolone (122.35%), Corticosterone (27.68%), 11-deoxycortisol (4%), Progesterone (3.64%), Prednisone (0.85%), Testosterone (0.12%) and ≤ 0.10%: Androstenedione, Cortisone and Estradiol. (<http://www.enzolifesciences.com/ADI-900-071/cortisol-elisa-kit/>).

The method followed the kit instructions. The seal urine was defrosted to room temperature, vortex mixed on a WM20 Vortex Mixer and 1 ml transferred to a 1.5 m Eppendorf tube. Samples were centrifuged to remove suspended material at 18,000 rpm (38,750g) for 10 min.

Because the concentration range of CL in harbour seal urine was not known, there may have been too much cortisol or too little to meet the range of the ELISA kit (156-10,000 pg/mL cortisol, with a middle calibrator of 1,250 pg/mL cortisol). A test run was therefore undertaken using the creatinine (Cr) values to estimate the dilutions needed for the ELISA run of the harbour seal pup urine. The urine samples were diluted by various factors in de-ionised water, followed by 5 times dilution in the assay buffer to get a cortisol value close to the middle calibrator of the kit where the calibration curve is the most reliable.

The colour change in the samples was read using an ASYS Expert Plus UV Microplate, set to measure absorbance at 405 nm with background correction at 620 nm. A 4-parameter logistic curve fitting program was used to obtain the calibration curve. Original urinary cortisol concentrations in pg/ml were obtained by multiplying by the dilution factors in assay buffer and de-ionised water. The CL concentrations (pg/ml) in the original sample were obtained by multiplying by the dilution factors, and converted to mM/L using the molecular mass of cortisol. These values were then divided by mM creatinine to obtain the cortisol/creatinine (CL/Cr) molar ratio and thence converted to ng/mg Cr.

*ELISA results and comparison with results from UPLC-MS/MS analyses of duplicate samples*

The Cortisol concentrations from the 17 urine samples from the pups in Centre I analysed by ELISA ranged from 46 to 1164 ng/mg Cr, (median 99 ng/mg Cr). Eleven of these 17 samples were analysed for CL by both UPLC-MS/MS and ELISA. The correlation (Spearman coefficient r_s_) between the MS/MS and ELISA results was 0.417. However, since the upper critical value for n=11 is 0.536, the correlation was not significant. The ELISA result indicated a higher concentration for seven of the 11 samples, and the mean concentrations for the ELISA results (30.54, SD 14.24 ng/mg Cr) were 11% higher on average than for MS/MS (27.08, SD 16.89 ng/mg Cr).

| CL - ELISA ng/mg Cr | CL - UPLC - MS/MS ng/mg Cr |
| --- | --- |
| 14.48 | 6.78 |
| 30.17 | 18.66 |
| 21.12 | 10.48 |
| 40.13 | 23.2 |
| 42.11 | 26.26 |
| 30.45 | 33.21 |
| 19.5 | 18.09 |
| 22.23 | 19.79 |
| 31.45 | 34.5 |
| 19.83 | 38.24 |
| 64.44 | 68.63 |

Fig. 2.1. Cortisol (CL) concentrations (ng/mg Cr) comparing ELISA with subsequent UPLC-MS/MS analyses for 11 duplicate samples from centre 1 (2012).

From this comparison of ELISA and UPLC-MS/MS results for duplicate samples, it appeared that the ELISA results tended to indicate slightly higher cortisol concentrations than the UPLC-MS/MS, although the direction of the difference between the two sets of results was not consistent. It is likely, nevertheless, that the ELISA results were affected by other glucocorticoids in the urine sample. At the time of the ELISA analyses (2012) the finding from the UPLC/MS/MS analyses of later samples (2014–2016) of significant levels of endogenous prednisolone in the urine of rehab seal pups (see main text) was neither known nor predicted. The high cross-reactivity with prednisolone (PL) of 122.35%, was not considered by the ELISA kit manufacturer to be an issue for CL determination, since prednisolone was not thought to be naturally occurring in humans or other mammals in any other than trace amounts (Fidan et al. 2013, Famele et al. 2015). The present UPLC-MS/MS study of seal pup urine has indicated that – at least in seal pups – an immunoassay targeting cortisol may not give accurate concentrations of cortisol alone.

*References*

Fidan M, Gamberini, MC, Pompa G, Mungiguerra F, Casati A, Arioli, F. 2013. Presence of endogenous prednisolone in human urine. Steroids 78: 121–126.

Famele M, Ferranti C, Palleschi L, Abenavoli C, Fidente RM, Pezzolato M et al. 2015. Quantification of natural and synthetic glucocorticoids in calf urine following different growth-promoting prednisolone treatments. Steroids 104: 196–202.

Novak, MA, Hamel, AF, Kelly BJ, Dettmer AM and Meyer JS. Stress, the HPA axis, and non-human primate well-being: a review. Appl Anim Behav Sci. 2013; 143: 135-149.

Taussky, H.H., Kurzmann, R., 1954. A microcolorimetric determination of creatinine in urine by the Jaffe reaction. J. Biol. Chem. 208, 853–862.
